## Supplementary File 1 for "A Multi-center Cross-platform Single-cell RNA Sequencing Reference Dataset"

### Towards Best Practice in Single-cell Sequencing:

### A Comprehensive Multi-Center Cross-platform Benchmarking Study of Single-cell RNA Sequencing Using Reference Samples

Wanqiu Chen[1#], Yongmei Zhao[2,8#], Xin Chen[1,3#], Zhaowei Yang[4,1#], Xiaojiang Xu[5], Yingtao Bi[6], Vicky Chen[2,8], Jing Li[4], Hannah Choi[1], Ben Ernest[7], Bao Tran[8], Monika Mehta[8], Parimal Kumar[8], Andrew Farmer[9], Alain Mir[9], Urvashi Mehra[7], Jian-Liang Li[5], Malcolm Moos Jr.[10], Wenming Xiao[11]*, Charles Wang[3,1]*

[1]Center for Genomics, School of Medicine, Loma Linda University, 11021 Campus St., Loma Linda, CA 92350.

[2]CCR-SF Bioinformatics Group, Advanced Biomedical and Computational Sciences, Biomedical Informatics and Data Science Directorate, Frederick National Laboratory for Cancer Research, 8560 Progress Drive, Frederick, MD 21701.

[3]Department of Basic Sciences, School of Medicine, Loma Linda University, 11021 Campus St., Loma Linda, CA 92350.

[4]Department of Allergy and Clinical Immunology, State Key Laboratory of Respiratory Disease, Guangzhou Institute of Respiratory Health, the First Affiliated Hospital of Guangzhou Medical University, Guangzhou, Guangdong, 510182, P. R. China.

[5]Integrative Bioinformatics Group, National Institute of Environment Health Sciences, NIH, Department of Health and Human Services, Research Triangle Park, NC 27709.

[6]Abbvie Cambridge Research Center, 200 Sidney Street, Cambridge, MA 02139.

[7]Digicon Corporation, 7926 Jones Branch Drive, Suite 615, McLean, VA 22102.

[8]Sequencing Facility, Leidos Biomedical Research, Inc., Frederick National Laboratory for Cancer Research, 8560 Progress Drive, Frederick, MD 21701.

[9]Takara Bio USA, Inc., Mountain View, CA 94043.

[10]Center for Biologics Evaluation and Research & Division of Cellular and Gene Therapies, U.S. Food and Drug Administration, 10903 New Hampshire Avenue, Silver Spring, MD 20993.

[11]The Center for Devices and Radiological Health, U.S. Food and Drug Administration, FDA, Silver Spring, MD, 20993.

[#]Equal contribution (Co-1[st] author)

*Corresponding should be addressed to: CW and WX

#### Abstract

The task of integrating diverse single-cell RNA sequencing (scRNA-seq) datasets remains a major challenge for the field. There is therefore an urgent need for guidance in choosing algorithms that lead to accurate biological interpretations of varied data types acquired with different platforms. Using two well-characterized cellular reference samples, captured either separately or in mixtures, we compared different scRNA-seq platforms and several pre-processing, normalization, and batch-effect correction methods at multiple centers. Although pre-processing and normalization contributed to variability in gene detection and cell identification, batch-effect correction was by far the most important factor in correctly classifying the cells. Moreover, scRNA-seq dataset characteristics (e.g., sample/cellular heterogeneity and platform used) were crucial in determining the optimal bioinformatic method (e.g., Seurat v3 overcorrected batch-effects between dissimilar samples). However, cross-center/platform reproducibility was high when appropriate software/bioinformatic methods were applied. Our findings offer practical guidance for optimizing platform and software selection when designing a scRNA-seq study.

#### Introduction

Single-cell RNA sequencing (scRNA-seq) allows transcriptomic profiling of individual cells in unprecedented detail[1-5], prompting increasingly widespread application of this technology. However, investigators seeking to adopt this technology are presented with a bewildering choice of analytical platforms and bioinformatics methods, each with its own set of capabilities, limitations, and costs[1-3,6, 7,8, 9,10, 11].

Various aspects of this problem have been examined recently[12,13,14]. Investigators from the Human Cell Atlas (HCA) consortium performed a comprehensive multi-center study that compared 13 different scRNA-seq protocols using a reference sample containing cells from human, mouse, and dog[15]. Consistent with the previous reports[12,13,14], this group observed not only striking differences among protocols in quantifying gene expression and identifying cell-type markers, but also found large cross-protocol differences in their capacity to be integrated into reference tissue atlases[15]. The large number of methods compared, and the diversity of cells analyzed will likely establish this work as an important milestone for the field; however, comparison of data analysis methods was not a major emphasis of this work. Tian et al. conducted a single-laboratory comparison of three batch-correction methods applied to four scRNA-seq datasets using mixtures of five lung cancer cell lines as reference material[16]. These investigators found significant differences between the methods tested; however, the best-performing of the methods evaluated nevertheless identified six clusters of cells in a mixture consisting of five cell lines. Recently, Tran et al. assessed 14 bioinformatic methods using the datasets from several public domain sources to simulate five different cellular input scenarios[17]; this group nevertheless did not compare various data preprocessing or normalization procedures and evaluated comparatively simple sample composition scenarios, i.e., most consisting of only two batches.

These studies used mixtures of cells exclusively, making it difficult to distinguish biological variability between heterogenous cell types (cell classification) from purely technical factors (analytical technology platform, institutional or other interlaboratory differences in cell handling, library protocols, and data processing methods; i.e., batch correction). This ambiguity makes it difficult to identify the various factors that affect the accuracy of biological classification of the cells analyzed. Therefore, there is no systemic multi-center study that evaluates the influence of technology platform, sample composition, and bioinformatic methods (including preprocessing, normalization, and batch-effect correction) using publicly available standard reference samples and datasets consisting of both mixed and non-mixed biologically distinct samples.

As part of the 2nd phase of the Sequencing Quality Control (SEQC-2) Consortium, we designed a comprehensive multi-center study to meet this need, anchored on two well-characterized, biologically distinct commercially available reference cell lines[18] for which a large amount of multi-platform whole-genome sequence (WGS) has been obtained (see the companion manuscript accepted by *NBT*)**.** Our study compared a breast cancer cell line vs. a “normal” B lymphocyte cell line to model practical, realistic situations in which malignant and normal tissues are analyzed in parallel for diagnostic purposes or for designing personalized medicine therapies[19]. In total, 20 scRNA-seq datasets derived from the two cell lines, analyzed either separately or in mixtures, were generated using four scRNA-seq platforms across four centers.

We compared six scRNA-seq data preprocessing pipelines, eight normalization methods, and seven batch correction algorithms. Our analyses indicated that although pre-processing and normalization contributed to variability in gene detection and cell classification, batch effects were quite large, and the ability to assign the cell types correctly across platforms and sites was dependent on the bioinformatic pipelines, particularly the batch correction algorithms used. In many scenarios, Seurat v3, Harmony, BBKNN, and fastMNN all corrected the batch effects fairly well for scRNA-seq data derived from either biologically identical or dissimilar samples across platforms and sites. However, when samples containing large fractions of biologically distinct cell types were compared, Seurat v3 over-corrected the batch-effect and misclassified the cell types (i.e., breast cancer cells and B lymphocytes clustered together), while limma and ComBat failed to remove batch effects. Overall, cross-center/platform consistency was high when appropriate bioinformatic methods were applied. These large cross-platform/site scRNA-seq datasets using well-characterized and banked cell lines will provide not only a valuable resource for biomedical research, but also represent a useful resource for benchmarking single-cell technologies and bioinformatics pipelines for the single-cell sequencing community, and for integrating diverse datasets contributed to large collaborative projects such as the HCA. Our findings offer practical guidance for selecting the combination of technology platform and bioinformatic methods best suited to the scientific question being addressed.

#### Results

##### 1. Study design

We used four scRNA-seq platforms: 10X Genomics Chromium, Fluidigm C1, Fluidigm C1 HT, and Takara Bio ICELL8, across four sites (LLU, NCI, FDA, and Takara Bio), using two well-characterized reference cell lines[18], a human breast cancer cell line (sample A) and a matched control “normal” B lymphocyte cell line (sample B) derived from the same donor (**Figure 1a)**. Overall, we generated 20 different scRNA-seq datasets including four different 3’-transcript scRNA-seq datasets, 10X_LLU, 10X_NCI, 10X_NCI_M (modified shorter sequencing protocol), C1_FDA_HT, and three different full-length transcript scRNA-seq datasets, C1_LLU, ICELL8_PE, and ICELL8_SE (**Table 1**). For the 10X single-cell platform, we compared the standard sequencing protocol (26x98 bp) with the modified sequencing method (26x57 bp) using the same scRNA-seq libraries. For the ICELL8 platform, we also compared paired-end (75x2 bp) with single-end (150 bp) analysis. We applied three different pre-processing pipelines either for the 3’-transcript scRNA-seq or for the full-length scRNA-seq (**Suppl. Table 1 & 2**). We also evaluated eight different normalization methods and seven different batch-effect correction algorithms as well as the consistency of scRNA-seq across sites/platforms with bulk cell RNA-seq on the two cell lines (triplicates each, six RNA-seq datasets).

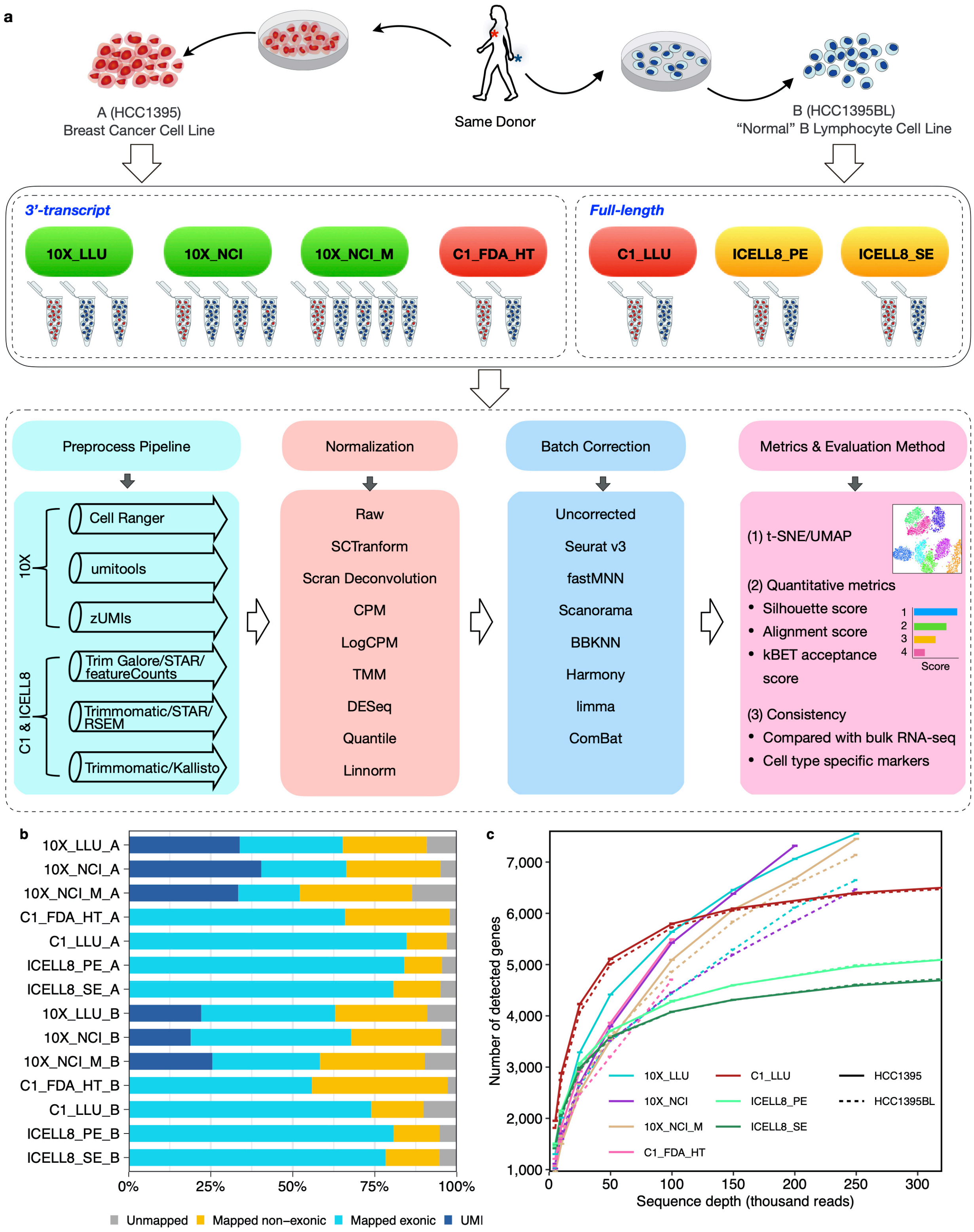


**Figure 1. Overall study design, scRNA-seq mapping, and numbers of genes detected across platforms.**
**(a)** Schematic overview of the study design (see detailed descriptions and notations in the Methods). Two reference cell lines (sample A & sample B) were used to generate scRNA-seq data across four platforms (10X Genomics, Fluidigm C1, Fluidigm C1 HT, and Takara Bio ICELL8), four testing sites (LLU, NCI, FDA, and Takara Bio). At the LLU and NCI sites (10X), mixed single-cell captures and library

constructions were also prepared with either 10% or 5% cancer cells spiked into the B lymphocytes. At the NCI site, single-cell captures and library constructions were also performed with methanol-fixed cell mixtures (5% cancer cell spiked into B lymphocytes, Fixed 1 & 2). One set of 10X scRNA libraries from NCI was also sequenced using a shorter modified sequencing method. Bulk cell RNA-seq was also obtained from these cell lines, each in triplicate. See Methods for details about study design. **(b)** For both the breast cancer cell line (A) and the B lymphocyte cell line (B) across 14 pair-wise datasets, percentage of reads mapped to the exonic region (blue), non-exonic region (orange), or not mapped to the human genome (gray). For Unique Molecular Identifier (UMI) methods (10X), dark blue indicates the exonic reads with UMIs. **(c)** Median number of genes detected per cell at different sequencing read depths. Solid line represents the breast cancer cell line (A). Dashed line represents the B lymphocyte cell line (B).

The overall assessments of the data generation and data QC are in the Online Methods section and in **Supplementary Table 1** and **Supplementary Figure 1.** We obtained twenty different scRNA-seq datasets from a total of **31,887** single cells with either 3' or full-length scRNA-seq data (**Suppl. Table 1 & Suppl. Fig. 1**).

To investigate the effect of sequence depth on the number of genes detected and saturation rates across all platforms and scRNA-seq datasets, we down-sampled the different datasets to varying read depths (**Fig. 1c & Suppl. Fig. 3**). We observed that the number of genes detected per cell increased rapidly with sequencing depth per single cell up to 100k reads/cell for both cancer cells (A) and B-lymphocytes (B), particularly much faster for the C1 before 50k read-depth. With increasing depth, gene counts ultimately plateau, as expected. However, the rate of saturation was slower after 100k reads for the full-length technologies (C1_LLU and ICELL8), with fewer additional genes being detected for the same increase in sequencing depth when compared with 3' scRNA-seq technologies (**Fig. 1c & Suppl. Fig. 3**). One interpretation of this is the higher complexity of full-length libraries due to sampling fragments across a gene's entire transcript as compared to 3' technologies which are biased to sampling only the end of the gene by design. A caveat here is that in the case of the 3' technologies, few cells within the population obtain this read depth (**Suppl. Fig. 1**), so that on a population level, the full-length technologies provide greater sensitivity. Our data confirms this observation and libraries from full-length technologies have higher library complexity and provide better representations of the captured transcripts with lower sequencing depth than 3' based technologies. However, the continuous increase in the number of genes detected with deeper sequencing observed for the 10X scRNA-seq data may be dependent on the transcript content of the cell-type; dependence of saturation rate on cell RNA content has been reported previously (https://kb.10xgenomics.com/hc/en-us/articles/115005062366-What-issequencing-saturation).

For benchmarking scRNA-seq data, we also identified a large number of differentially expressed genes (DEGs) between the two cell lines (six RNA-seq datasets, triplicate in each cell line) at the population level (**Suppl. Data 1**, using fold-change ≥ 2 plus P-value ≤ 0.01, FDR= 0.05). The bulk cell RNA-seq sequencing depth and mapping QC are shown in **Supplemental Table 4** and **Supplemental Fig. 4**.

**2. Effects of data pre-processing**

For the UMI based scRNA-seq data, we compared three pipelines for pre-processing the data: Cell Ranger 3.1 (10X Genomics)[20], UMI-tools[21], and zUMIs[22], and examined the consistency between the three pipelines regarding the number of barcoded cells identified and the number of genes detected per cell (**Fig. 2a & 2b,**

**Suppl. Table 2**). For the non-UMI based scRNA-seq data, we examined three additional pre-processing pipelines: FeatureCounts[23], Kallisto[24], and RSEM[25] (**Fig. 2d & Suppl. Table 3**), which included trimming processes (cutadapt[26] or trimmomatic[27]), alignment (STAR[28] and Kallisto), and gene counting (FeatureCounts, Kallisto, and RSEM). For simplicity, we used FeatureCounts, Kallisto, and RSEM for the non-UMI-based platforms. We observed that, for the UMI-based scRNA-seq data, there were variations across the three pipelines both in the number of cells identified and number of genes detected per cell (**Fig. 2a & 2b**). Cell Ranger v3 was the most sensitive method for cell barcode identification. Umitools and zUMIs filtered most low gene/transcript expressing cells, but detected more genes per cell. Nonetheless, the gene expression level and the consensus genes per cell were highly correlated between any two of the UMI-based pre-processing pipelines; with Umi-tools and zUMIs showing the highest concordance (**Fig. 2c**).

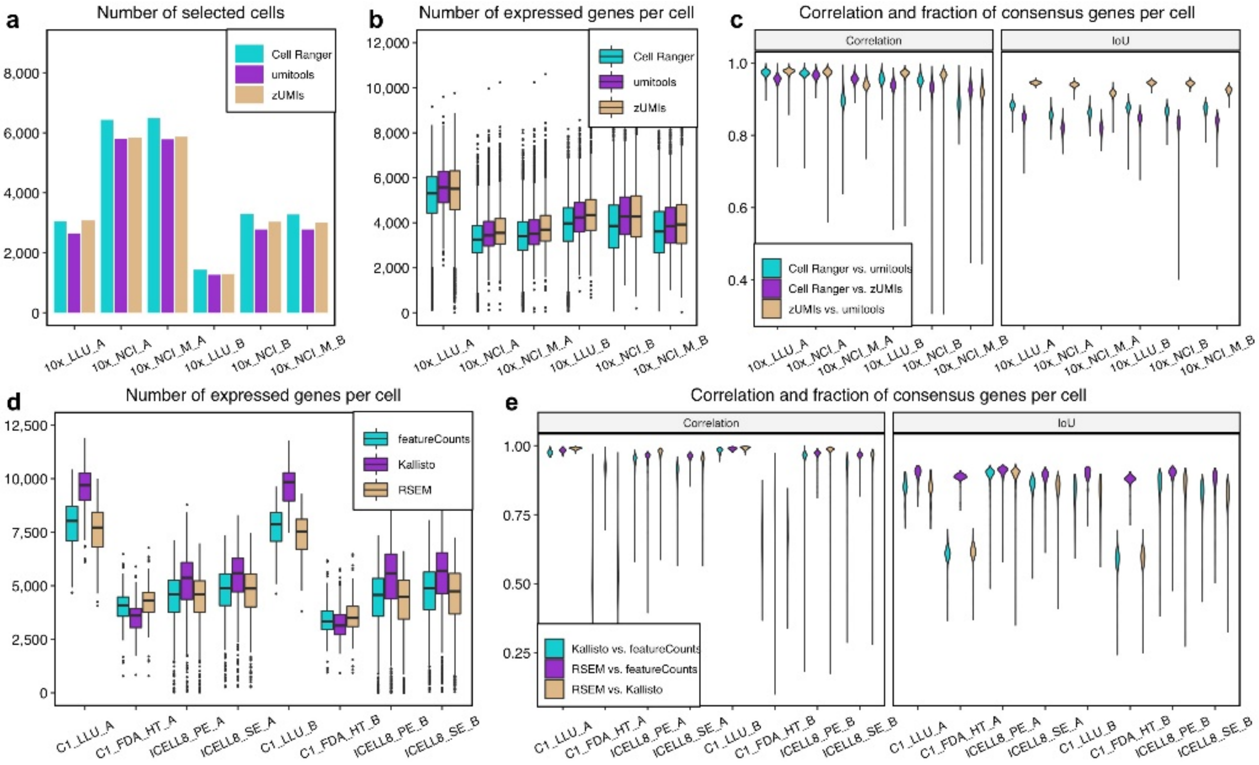


**Figure 2. Effect of pre-processing pipeline on the number of genes detected with UMI- and non-UMI-based scRNA-seq datasets.**

**(a-c)** Evaluation of the UMI-based (10X) data with Cell Ranger, UMI-Tools, or zUMIs. **(d-e)** Evaluation of data from non-UMI based technologies C1 full transcript, C1 HT, and ICELL8 full-length transcript using FeatureCounts, Kallisto, or RSEM. **(a)** Bar plot showing the number of cells captured with UMI-based technology. **(b)** and **(d)** Box plot showing the number of genes detected per cell in UMI-based and non-UMI based technologies. **(c)** and **(e)** Violin plots showing the gene expression correlation and consensus genes [represented by IoU (Intersection over Union)] per cell between any two pipelines in UMI-based and non-UMI based technologies.

For non-UMI based scRNA-seq data, much larger variation was observed in the number of genes detected across the three different pre-processing pipelines (**Fig. 2d**). Interestingly, we found that Kallisto identified a significantly higher number of genes per cell in the full-length transcript scRNA-seq datasets (C1_LLU and ICELL8) and the fewest genes per cell in the C1-FDA_HT (3' counting) datasets (**Fig. 2d**). In addition, the consensus genes per cell from the Kallisto pipeline differed significantly from the gene list generated by the other two pipelines for the C1-HT 3' method. This suggests that the performance of genome alignment-based (RSEM and FeatureCounts) and pseudo-aligner tools (Kallisto), which are most commonly used for full-length isoform analysis algorithms, might underperform when pre-processing scRNA-seq data from 3'-based technologies. Overall, we found that the gene expression (counts) and the fraction of consensus genes per cell were highly variable across three pre-processing pipelines, both for UMI- and non-UMI based scRNA-seq datasets (**Fig. 2c & 2e**). To simplify our comparison, we used Cell Ranger, one of the most popular UMI-based methods for 3' count technologies, and FeaturesCounts for non-UMI based technologies, since it was more consistent with RSEM than Kallisto, in all of our subsequent analyses.

##### 3. Effects of normalization

scRNA-seq data characteristically demonstrate substantial numbers of zero read counts[29]. This can be due to both biological (e.g., bi-stable gene regulation) and technical reasons (e.g., 'drop out' due to Poisson sampling limitations or limited efficiency of reverse transcription), making the normalization of scRNA-seq data very challenging. So far, global scaling normalization methods developed for bulk cell RNA-seq data have been used fairly often for scRNA-seq data including CPM (Counts per Million), UQ (upper quantile), TMM (trimmed mean of M-value), and DESeq[30]. However they were never systemically evaluated using a standard reference scRNA-seq dataset until Tian's computational integration analysis, which involved two batches of mixed sample sets, but no non-mixed samples captured independently[16]. Regression-based methods have also been proposed to remove known nuisance factors in scRNA-seq data. There are also methods specifically tailored to scRNA-seq datasets, such as SCTransform[31], scran[32], SCnorm[33], and Linnorm[34] etc.. SCTransform was developed most recently and has been integrated within Seurat v3[35]. Nevertheless, a systematic, thorough evaluation of these methods using scRNA-seq datasets derived from standard reference samples analyzed at multiple centers using multiple platforms is very much needed by the community.

We evaluated eight different normalization methods including SCTransform, Scran deconvolution[36], CPM, LogCPM, TMM, DESeq, Quantile, and Linnorm, using the silhouette width metric, which evaluates how well the two samples from the same cell type are grouped with each other (**see Methods**). We noticed that TMM and Quantile failed to normalize either the breast cancer or B-lymphocyte samples, with silhouette scores that were similar to the un-normalized raw data (**Fig. 3a-g**). The other methods provided similar normalization as measured by silhouette score. SCTransform seemed to perform slightly better than Scran deconvolution, LogCPM, and Linnorm in that it had the least variation among all normalization methods (**Fig. 3a-g and Suppl. Fig. 5a-g)**. When comparing silhouette scores across different scRNA-seq platforms and datasets, we noticed that the 10X scRNA-seq data gave consistently lower scores than either the C1 data (both full-length and 3' across two sites) or the ICELL8 data (**Suppl. Fig. 5a-g**).

Distinguishing between irrelevant variations and biological changes of interest can be challenging. A common approach is to pre-process single-cell RNA-seq data to remove biological differences such as high mitochondrial gene levels in subpopulations or different cell cycle stages. When multiple weak and non-independent factors are present in a dataset, it is also common to apply computational strategies such as linear regression with read depth normalization, or tailored methods such as scLVM[37] to remove variations before applying subsequent analysis methods[37]. We found that regressing out mitochondrial genes did not improve the downstream clustering results (**Suppl. Fig. 6a-h**), and that regressing out the number of genes detected or using a sequencing depth approach did not improve silhouette scores (**Suppl. Fig. 7**).

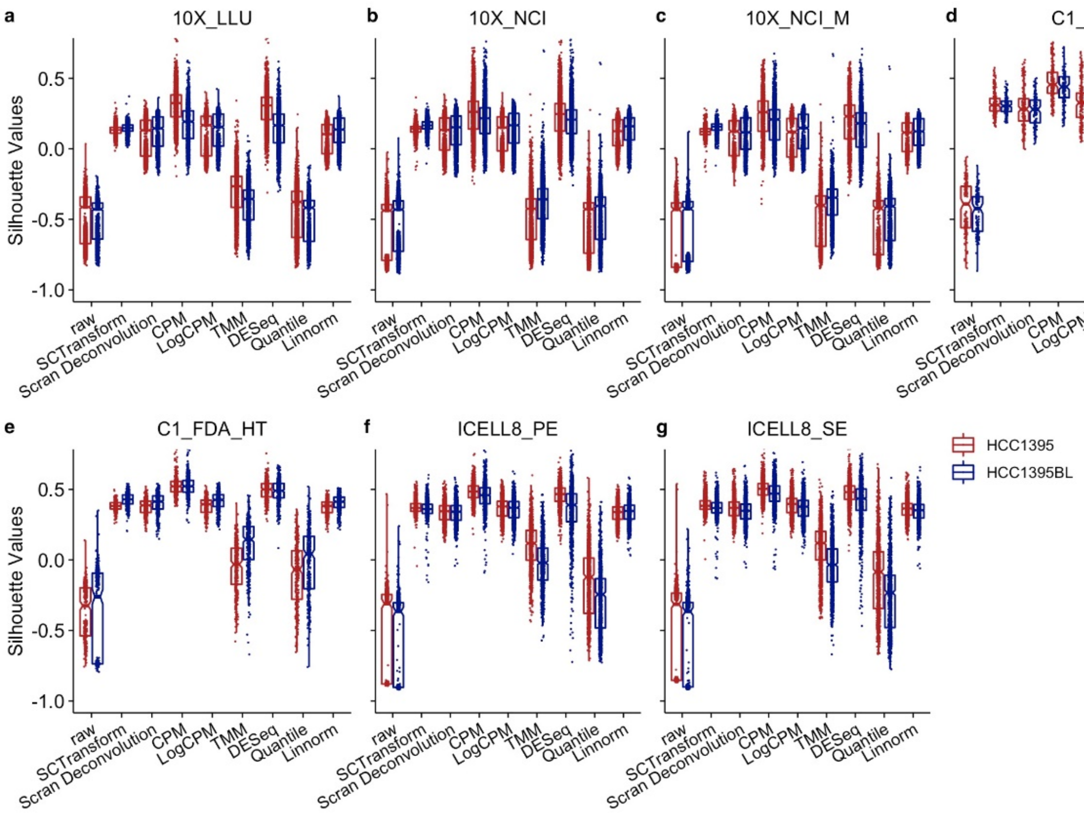


**Figure 3. Silhouette score boxplot comparing eight normalization methods.**
**(a-g)** Boxplot of silhouette values stratified by eight normalization methods across 14 datasets, including 10X_LLU (**a**), 10X_NCI (**b**), 10X_NCI_M (**c**), C1_FDA_HT (**d**), C1_LLU (**e**), ICELL8_PE (**f**), and ICELL8_SE (**g**) in breast cancer cells (HCC1395), and B lymphocytes (HCC1395BL). Eight normalization methods included SCTransform, Scran Deconvolution, CPM, LogCPM, TMM, DESeq, Quantile, and Linnorm. For each dataset, reads of each cell were down-sampled to two different read depths (10K and 100K per cell) before calculating the silhouette width values. LogCPM normalization performed fairly well and was used as the default normalization for our subsequent batch-effect correction benchmarking analyses. Two normalization methods developed for bulk cell RNA-seq had the lowest scores (TMM and Quantile).

Since log transformation has a high impact on downstream feature selection and clustering analysis and our analysis showed it performed fairly well, we used logCPM in our subsequent batch-effect and benchmarking evaluations except where specific normalization methods were embedded in some pipelines.

#### 4. Batch effects and batch effect corrections evaluated by t-SNE, UMAP, and quantitative metrics

As noted earlier, variability between data sets can result from both technical and biological factors[29, 38]. We benchmarked seven algorithms for batch-effect correction including Seurat v3[35], fastMNN or Mutual Nearest

Neighbors (MNN)[6], Scanorama[8], Batch-Balanced k-Nearest Neighbors (BBKNN)[9], Harmony[10], limma[39], and ComBat[40]. We visualized clustering projections with both t-SNE and UMAP[41]**,** and applied quantitative metrics including silhouette width, kBET[42], and alignment score[7] in four different sample scenarios to evaluate the batch-effect removal and cross-platform/center dataset integration as measured by clusterability (ability to separate dissimilar cell types) and mixability (ability to group similar cell types). kBET and alignment score quantify mixability, whereas silhouette width score, which was applied to our sample scenarios #1 and #4, quantifies how well two different types of cells are separated from each other.

First, we interrogated all 20 scRNA-seq datasets including the spike-in samples, taking the gene counts based on the preprocessing pipelines selected as above using either logCPM or the normalization method embedded in the pipelines (e.g., SCTransform for Seurat v3) (scenario #1-all datasets combined including mixed and non-mixed, with large proportions of two dissimilar types of cells). We sought to determine: 1) which of the algorithms could remove the batch-effects and also separate the two cell types correctly (clusterability/cell identification); and 2) how well cells of the same type from different batches were grouped together (mixability). There were large variations across platforms and centers (**Fig. 4a, Suppl. Figs. 8a, 9a-b, 10a, 11a, left panels, uncorrected**). However, BBKNN (ranked on the top at clusterability, **Fig. 4e**), fastMNN, and Harmony were effective in removing batch-effects and separating cancer cells from B cells. BBKNN performed well in grouping B cells together from different batches, but was the worst of the methods tested for cancer cells by this criterion-mixability (**Fig. 4a-b/e/g and Suppl. Fig. 12**). Seurat v3 was best at grouping similar cells from different batches together, but over-corrected and clustered B lymphocytes and breast cancer cells, two highly dissimilar cell types, together— a misclassification (**Fig. 4a/e/g, Suppl. Figs. 8a, 9a-b, 10a, 11a and 12**). Scanorama, limma, and ComBat were not only unable to remove batch effects but also failed to separate cancer cells from B cells (**Fig. 4a/e/g, Suppl. Figs. 8a, 9a-b, 10a, 11a and 12**). However, when only data from the 10X platform were analyzed, Scanorama both separated dissimilar cells clearly and grouped similar cells together very well, regardless of center (**Suppl. Figs. 13, 8d**, **10d**, **11d**, **Fig. 4d)**. For fastMNN, a spiked in sample was required to provide a subpopulation of cells common to all samples analyzed. We also found that the order of loading the datasets into fastMNN was critical for correcting batch effects; specifically, the mixed, or most heterogenous, sample should be loaded into the pipeline first (**Suppl. Figs. 14 & 15**).

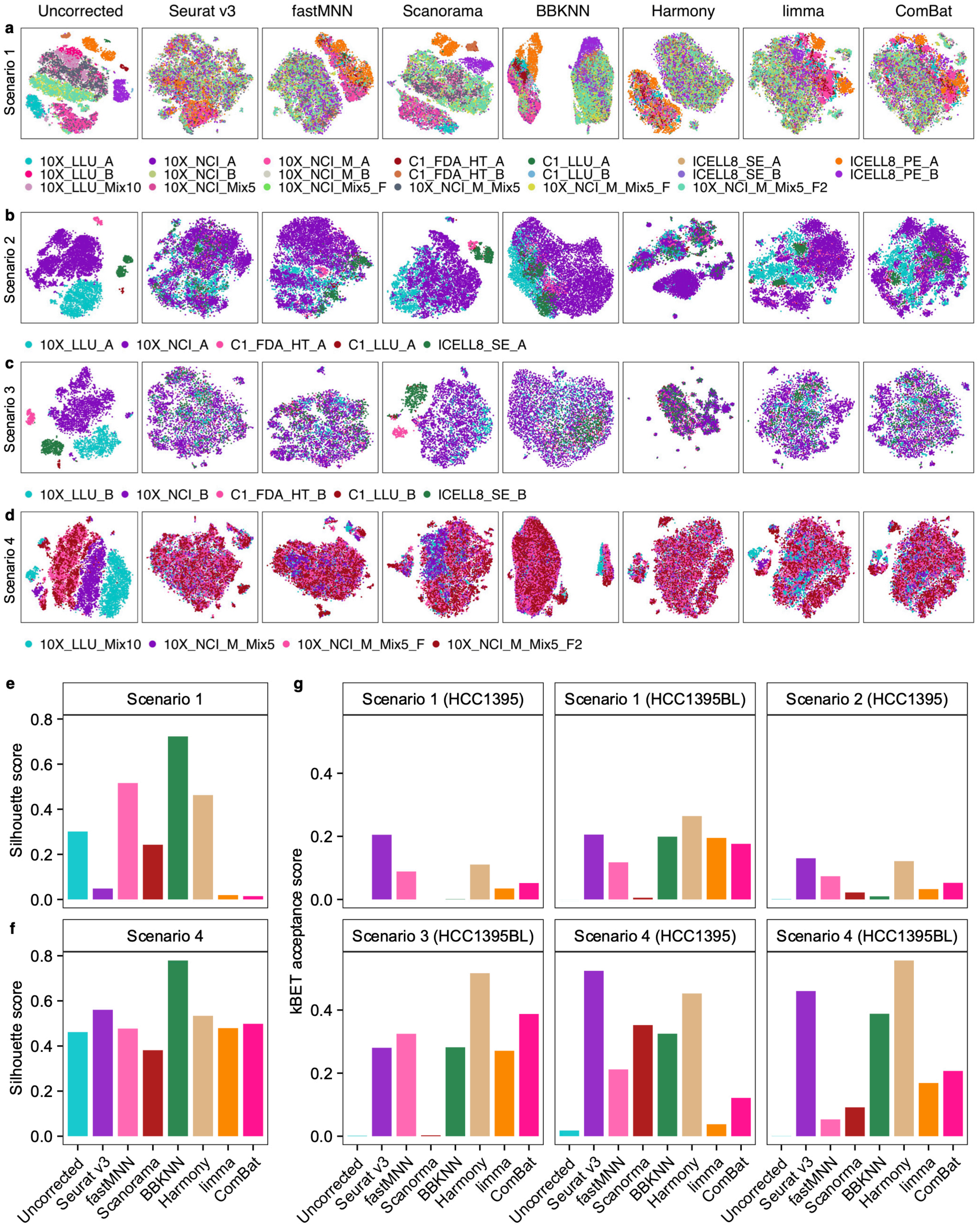


**Figure 4. Batch-effect corrections evaluated in four different sample composition scenarios.**
**(a)** Batch-effect corrections performed using 20 scRNA-seq datasets across all platforms and centers. Dataset is encoded by color. Samples from 10X were down-sampled to 1200 cells. **(b)**-**(c)** Batch-effect corrections performed using five batches of cross-platform/center scRNA-seq data obtained from either breast cancer cells **(b)** or B lymphocytes **(c)**. **(d)** Batch-effect corrections performed using four batches across two centers of scRNA-seq data derived from spiked-in mixtures of cells in which either 5% or 10% cancer cells

were spiked into the sample B cells. The top 2000 highly variable genes (HVGs) were used for the batch correction evaluation in all scenarios (**a-d**). All plots are t-SNE projections except for the BBKNN pipeline, into which UMAP is integrated and shown instead. (**e-f**) Silhouette width score quantifying the batch-effect on clusterability for the scenario #1 (**a**) or #4 (**d**), respectively. (**g**) kBET acceptance score calculated using the cross-platform/center scRNA-seq data acquired from the two cell lines separately (i.e., either breast cancer cells only or B-lymphocytes only) for all four scenarios (**a-d**, also labeled as #1-#4).

Second, we evaluated the batch-correction methods using scRNA-seq data from each cell line separately (scenarios #2 & #3; **Fig. 4b & c**). We also evaluated samples with 5% or 10% of cancer cells spiked into the B cell sample, and analyzed with the 10X Genomics platform across two centers in four different batches (scenario #4-mixtures; **Fig. 4d**). Without batch correction, t-SNE & UMAP showed that the cells were clustered separately by batch, and that similar cells were not evenly mixed, indicating large variations and/or strong batch effects (**Fig. 4b-d & Suppl. Figs. 8b-d, 10b-d/11b-d, left panels**).

In scenarios #2 and #3 (**Fig. 4b-c**), we observed that Harmony, Seurat v3, and fastMNN removed the batch effects fairly well for the breast cancer cells and B cells (**Fig. 4b-c/g, Suppl. Figs. 8b-c, 10b-c & 11b-c**). Consistent with our finding in scenario #1, BBKNN had the poorest mixability in cancer cells despite its good performance in grouping B cells together and batch-effect removal. In contrast, Scanorama performed poorly in both batch-effect removal and cellular mixability (**Fig. 4b-c/g, Suppl. 8b-c/9a-b & 12**). For B cells, which are relatively more homogeneous than the breast cancer cells, limma and ComBat also seemed to perform well in batch-effect removal and in grouping similar cells together (**Fig. 4c/g, Suppl. Figs. 8c, 10c & 11c**). The t-SNE plots in **Figure 4d** illustrate the performance of batch-correction methods applied to a scenario when a dataset was generated from mixtures containing a small number of distinct cells (scenario #4-mixtures). In these spike-in datasets, all methods were able to remove batch-effects and separate the spiked in cancer cells from B cells discretely, with BBKNN performing the best (**Fig. 4d/f**), followed by Harmony and Seurat v3 (notwithstanding that it failed in scenario #1), whereas Harmony, Seurat V3, BBKNN, and Scanorama all performed well in grouping similar cells together (**Fig. 4d/f-g, Suppl. Figs. 8d, 10d/11d and 12**).

We also compared mnnCorrect vs. fastMNN using the 20 scRNA-seq datasets, and found both versions performed similarly in t-SNE and UMAP projections. However, fastMNN took much less computation time: for example, 51.1 seconds for fastMNN vs. 62781.8 seconds for MNN scenario #1 (1,229-fold faster, **Suppl. Fig. 16**).

CellRanger 3.1 allows some cells with extremely low gene expression level to be identified. As a result, significantly more cells were detected using CellRanger 3.1 than with CellRanger 2.0 (**Suppl. Table 5**). We therefore compared batch correction results obtained with the two versions for all four sample combination scenarios. Overall, even though there was substantial consistency between CellRanger 3.1 and 2.0 preprocessed data; we noticed that the batch corrections performed better using CellRanger 2.0 (**Suppl. Figs. 17 & 18)**. As part of our cross validation, we also performed batch correction analysis on the four scRNA-seq datasets from Tian et al.[16], which were generated from mixtures of either 3 or 5 lung cancer cell lines in two batches. Consistent with the findings using our own datasets, our analysis using Tian et al. datasets showed that fastMNN, Seurat v3, and Harmony performed well (**Suppl. Figs. 19 & 20**), whereas in the Tian et al. analysis, both Seurat v2 CCA (Canonical Correlation Analysis) and MNN failed to separate cells from five different cell lines[16].

**5. Consistency of global gene expression across platforms and sites**

We first evaluated global gene expression cross-platform consistency using scatter plots and the common transcripts detected across seven scRNA-seq datasets for either sample A or B (**Suppl. Fig. 21a-b**). The bar chart plots clearly show that the bulk cell RNA-seq had a much wider range of genes with high expression than any of the scRNA-seq platforms where majority of genes have very low UMI counts. Overall, our scatter plot analyses showed cross-platform correlation coefficients between single-cell datasets of 0.8-0.98; correlation coefficients between scRNA-seq and bulk cell RNA-seq were lower, which could be due to the large differences in gene expression counting between bulk and scRNA-seq. Pearson correlation coefficient analysis indicated a higher intra-platform than inter-platform correlation for both sample A and sample B. For example, 10X scRNA-seq data had high correlations across centers, ≥ 0.94 for sample A and 0.98 for sample B.

We then evaluated global gene expression consistency across different platforms and sites by calculating a pairwise Pearson correlation (**R**) based on the percentage of cells (**see Methods**) that expressed 500 abundant, 500 intermediate, and 500 scarce genes, as defined by bulk cell RNA-seq data (**Suppl. Fig. 22a-f**). To account for variable sequencing depth across the different datasets, we performed down sampling to 100K reads for each dataset. At this read depth, we observed a much higher Pearson correlation when using the 500 highly-expressed genes than when using 500 intermediate or 500 scarce genes in both cell types (**Fig. 22a-f**). We also observed higher consistency between the sites using the same platform or type of technology (i.e., 10X, ICELL8, or C1). Even with the 500 low-abundance genes, we observed a reasonably good Pearson correlation between sites or within 10X 3', ICELL8, or C1 technologies (**Fig. 22c & 22f**). However, the consistency (Pearson correlation) within 3' technologies (10X and C1_FDA_HT) or within full-length (C1_LLU, ICELL8_PE, ICELL8_SE) platforms was not always better than that between 3' and full-length platforms. Nevertheless, we want to caution that there might be some biases in this analysis since the cell numbers were very different across platforms, i.e., only 66 or 80 single cells for the C1 full-length versus up to a few thousand cells for the 10X platform. Thus, the influence of variation due simply to sampling must be considered.

We further compared the single-cell gene expression profiles [log(CPM+1) with normalized counts] across four different classes of RNA including protein coding RNA, antisense RNA, lincRNA, and miscRNA (**Suppl. Fig. 23**). As a comparison, the bulk cell RNA-seq gene expression profile was also plotted side-by-side. We noticed that ICELL8_SE gene expression profiles showed relatively higher detection sensitivity for the lower abundance transcripts. The 10X technology also seemed to show good detection sensitivity in the lower abundance transcripts for the protein coding RNA, antisense RNA, and lincRNA and there was high consistency across three 10X scRNA-seq datasets (10X_LLU vs. 10X_NCI vs. 10X_NCI_M). Gene counts across all scRNA-seq platforms and datasets for the protein coding RNAs were comparable. Interestingly, for the C1 platforms (full-length and 3'), the detection range was compressed, with much lower log(CPM+1) values for antisense RNA and lincRNA.

#### 6. Consistency of cell-type specific markers across scRNA-seq platforms and sites

We exploited feature plotting using the top ten B cell-specific DEGs[43, 44] and top ten cancer specific DEGs based on the DEGs derived from the bulk cell RNA-seq to further evaluate single-cell gene expression consistency prior to and post fastMNN correction across all scRNA-seq datasets and platforms. Clearly, prior to fastMNN batch-effect correction, neither of the two cell types were clustered together, and there was no clear separation between them (**Fig. 5a & 5c**). However, after applying fastMNN, cells expressing B-cell specific vs. breast cancer specific marker genes were nicely clustered together and there was clear separation between the two cell types (**Fig. 5b & 5d**).

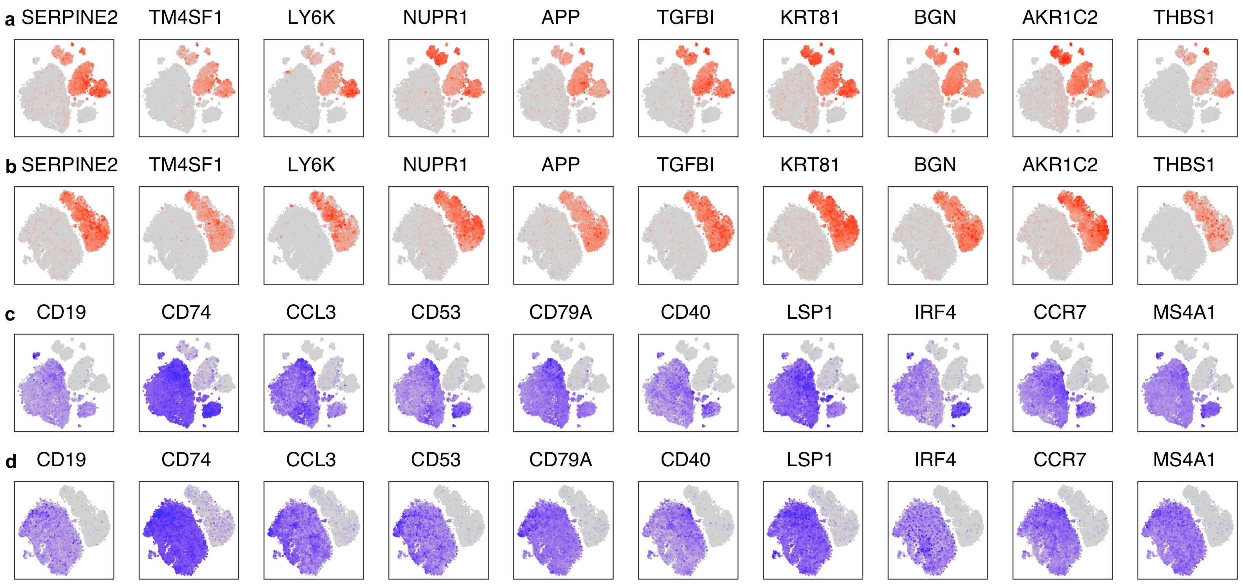


**Figure 5. Feature plots showing cell type clustering based on cell type-specific marker genes across 20 scRNA-seq datasets.**

Feature plots generated using the top 10 breast cancer (**a-b**) or B lymphocyte (**c-d**) specific DEGs across 20 scRNA-seq datasets before (**a & c**) and after (**b & d**) fastMNN correction. Genes with relatively high expression in each cell were highlighted in brick red (sample A) or blue (sample B) in t-SNE plots.

We further compared the consistency of single-cell gene expression across platforms for CD40, CD74, and TPM1 with a subsampling at 100k reads for each dataset. The rationale for selecting these three markers to benchmark the transcript detection consistency across platforms was based on both their cell-type specificity and their expression levels, either intermediate or highly abundant. The B-cell specific marker gene CD40 was most often expressed at an intermediate level (1 ≤ CPM < 10) per cell, and it was detected in as few as 24.9% of cells with the C1_FDA_HT to as many as 53% with the C1_LLU. A significant percentage of cells (44% to 44.6% for 10X and 23.2% to 28.1% for C1 and ICELL8) expressed this gene at levels close to the limit of detection (CPM < 1). In contrast, the CD40 transcript was detected at either low or near noise levels (CPM <1) in breast cancer cells (**Suppl. Table 6**). However, CD74, also a B-cell specific marker gene, was much more abundant (CPM ≥ 10) in almost all single B cells (98.9% - 100%) with excellent consistency across all platforms except for C1_FDA_HT, where 5% of the B cells had an intermediate level (1 ≤ CPM < 10, **Suppl. Table 7**). In contrast, CD74 was present at low or near noise levels (CPM < 1) in breast cancer cells. For this marker, full-length transcript technologies were more sensitive than the 3'-single-cell technologies (**Suppl. Table 7**). With some variation across platforms, a high percentage of single cells expressed TPM1 in breast cancer cells, but the detection level fell mostly within an intermediate level (1 ≤ CPM < 10). In B cells, consistent with their biological nature, there was little or no detection (CPM <1) (**Suppl. Table 8**).

#### Discussion

The availability of scRNA-seq datasets based on sustainable, well characterized reference samples that were processed across multiple platforms and centers is critical for benchmarking single-cell technologies and bioinformatic methods. Here, we benchmarked scRNA-seq performance across several popular platforms at multiple centers, focusing on the effects of bioinformatic processing; including preprocessing, normalization, and batch-effect correction. We analyzed two biologically distinct reference cell lines[18], either separately or as mixtures, for which a large amount of multi-platform WGS data are available. Our benchmark study has produced well-characterized reference materials (reference samples, datasets) and methods. In this regard, it will have similar resource value and utility for the single-cell sequencing community as the Zook at al. study[45], carried out by the Genome in a Bottle Consortium (GIAB), which aimed at developing reference materials, data and methods to enable translation of genome sequencing to clinical practice.

The findings from our study offer practical guidance for optimizing and benchmarking a platform or a protocol, and for selecting appropriate bioinformatics methods when designing scRNA-seq experiments. In our study, samples of both lines were distributed to different centers and grown out separately at these locations to reflect the sort of experimental variability likely to be encountered in real-world collaborations (in contrast to the situation with Genome-in-a-bottle or our companion manuscripts, in which identical aliquots of gDNA reference material were distributed to the study sites). As expected, site-to-site and platform-to-platform variability was large, but when an appropriate combination of computational methods was chosen, these effects could be corrected. For benchmarking a newly developed scRNA-seq platform or protocol, or for quality control while starting a scRNA-seq experiment regardless of platform, we recommend including a mixed sample with 5-10% of the reference breast cancer cells spiked into the reference B-cell line sample. Single cells of the mixed samples may be processed in different batches, while non-mixed samples of the breast cancer cells and B-cells should be processed in the same batch. The cells obtained from the provider should first be expanded for cryopreservation in multiple aliquots. Any aliquot of cells may then be sub-cultured for few rounds without significantly affecting the accuracy of cell clustering or identification, as demonstrated in our study. We also recommend obtaining bulk cell RNA-seq in triplicate for both reference lines. Gene detection for breast cancer (TMP1) or B-cell (CD40 and CD74) specific markers can be compared with our reference data (**Suppl. Table 6, 7, 8 and Fig. 5**).

The acquired scRNA-seq data can be pre-processed using any of the preprocessing methods ranked in **Figure 6a**, as appropriate to the scRNA-seq technology employed, and any of the normalization methods except for

TMM and Quantile, which are not recommended. If desired, the scRNA-seq data obtained can be merged with our reference datasets from any of the four data composition scenarios (**Fig. 4**), and analyzed using our benchmarked reference methods using different batch-correction algorithms (see **Suppl. Fig. 25 and File 1**). Moreover, our scRNA-seq reference datasets and reference bioinformatics methods can be a valuable resource for developing or benchmarking new methods, and we recommend using all four different sample/data composition scenarios (**Fig. 4**) to gain a thorough performance evaluation.

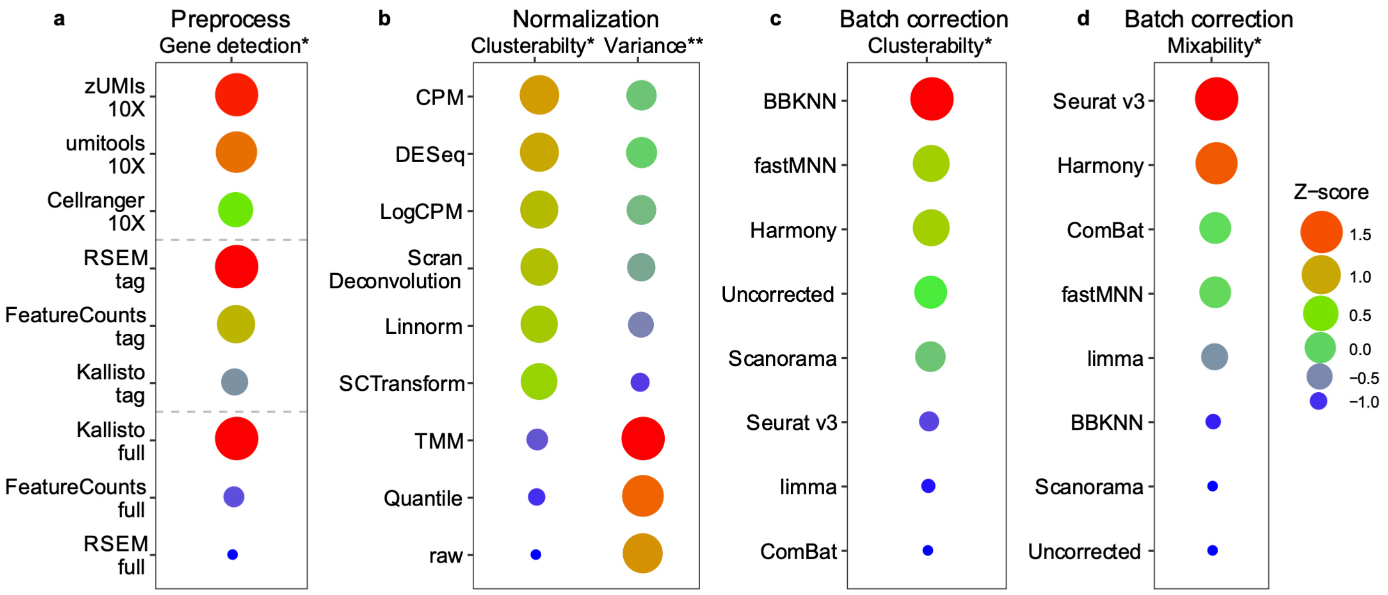


**Figure. 6. Performance ranking of bioinformatics metrics**

**(a)** Gene detection sensitivity measured separately for each of the three classes of scRNA-seq protocol: 10x-, non-10x-based 3' tagging, and full-length. **(b)** Normalization methods ranked by their clusterability as measured by Z-scores (either the median or the variance of the silhouette width across the 14 data sets). **(c)** Batch-correction methods ranked by their clusterability as measured by Z-score from the harmonic mean of the silhouette scores (scenarios #1 and #4). **(d)** Batch-correction methods ranked by their mixability as measured by Z-score from the harmonic mean of kBET acceptance scores (4 sample scenarios, #1 - #4). Z-scores are plotted as circles with their size and color scaled to the Z-score value from large to small, and red to blue. Note that larger Z-score values imply better performance, except for clusterability variance, where a smaller value is preferred. *Larger is better; **Smaller is better.

We found that pre-processing and normalization contributed to variability in gene detection and cell identification. For the UMI-based datasets, Cell Ranger v3 detected the most cells, while zUMIs detected the most genes per cell. For the non-UMI-based datasets, Kallisto identified the highest number of genes per cell in the full-length scRNA-seq datasets but the fewest genes per cell in the C1_HT dataset. Out of eight normalization methods evaluated, SCTransform, LogCPM, Scran deconvolution, and Linnorm all performed well, with the SCTransform displaying the lowest variance. In contrast, TMM and Quantile performed poorly across all datasets (**Figs. 2, 6**). Moreover, we found that regressing mitochondrial genes and normalizing UMI counts could not remove the batch effects (**Suppl. Figs. 6, 7**).

An ideal workflow would remove variability due purely to technical factors or sampling ambiguity without obscuring meaningful biological differences important to accurate classification of diverse cell types[15]. We reasoned that not all algorithms would perform equally well at these tasks, so we compared seven algorithms with respect to performance on two aspects of cross-platform data integration in addition to removing batch variations: clusterability and mixability. Seurat v3, Harmony, BBKNN, fastMNN, and Scanorama worked well in removing batch effects and classifying cells in some scenarios. However, sample heterogeneity, dataset composition, and platform used influenced the outcome of the analyses (**Fig. 4a-d, Suppl. Figs. 8a-d, 9a-b, 10a-d, 11a-d, 12-15**). For example, despite its high mixability for the data from the same cell line (**Fig. 4b-c/g**), Seurat v3 over-corrected batch effects and misclassified cells in datasets containing large fractions of highly dissimilar cells (**Fig. 4a/e, Suppl. Figs. 8a, 9a-b**). Scanorama worked well only for datasets generated entirely with the 10X platform (**Suppl. Fig. 13**), a caveat missed by the original developers of the algorithm[8], which indeed failed to remove batch effects from non-10X platform data, as confirmed by our re-analysis using their testing datasets (**Suppl. Fig. 24**).

BBKNN performed the best while limma and ComBat were poorest in cross-platform/center separation of two types of cells from each other, particularly when there were large proportions of dissimilar cells in the datasets (**Figs. 4a/e, 6c**). In contrast, Seurat v3, despite its failure to separate distinct cell types in the multi-platform datasets consisting of large proportions of each cell-type (**Fig. 4a/e**), worked well in the cross-center datasets comprising a large proportion one cell type spiked with a small portion of different cells (**Fig. 4d/f**). This method may be best suited to situations where the primary objective is to remove differences between datasets due mainly to technical sources of variations rather than to integrate data from biologically dissimilar cell populations. Consistent with this notion, Seurat v3, fastMNN, and Harmony all performed well in mixability for the scRNA-seq data derived from biologically identical or similar samples across platforms/sites, whereas limma and ComBat could mix B-cells well across platforms/sites, perhaps because these cells are more homogeneous (**Fig. 4b-c/4g, Suppl. Figs. 8b-c, 10b-c**). Indeed, for all algorithms, we found that the cross-platform/center mixability was better in the B-cells than in the more heterogeneous cancer cells (**Fig. 4b-c/4g**). However, for fastMNN/MNN, both the requirement for mixed samples and the order of importing data into the pipeline (i.e., the mixed samples should be imported first) were critical for effective batch-effect correction (**Suppl. Figs. 14, 15**). The presence of mixed cell types to provide a shared subpopulation across the batches to be corrected is consistent with the logic of MNN algorithm[6]. Thus, our findings clearly highlight that the choice of appropriate batch-effect method depends on the characteristics of the samples (e.g., heterogeneity, cell composition) and datasets (multiple vs. fewer platforms used).

Some combinations of different preprocessing methods with different batch-effect algorithms might not always work well. For example, Tian et al. examined imputation methods by combining Linnorm and Drimpute with MNN; TMM and Drimpute with Scanorama; and SAVER with Seurat using data from a mixture of five cell lines[16]. However, this approach failed to correct batch effects not only for Seurat and Scanorama, but also for MNN, the top performing method of those evaluated, which identified six clusters in the samples, instead of five cell lines (see their **Suppl. Fig. 8d**)[16]. However, when we applied our preferred preprocessing method to Tian's datasets, fastMNN, Seurat v3, and Harmony all performed really well in separating the five different cell lines (**Suppl. Figs. 19 & 20**).

Previous studies using mixtures containing heterogeneous distinct cell types have provided useful insights into bioinformatics methods particularly with regards to batch-effect correction algorithms[16]. One example is the requirement for shared subpopulations (anchor cells) between different datasets for Seurat and fastMNN, in order to integrate data from biologically dissimilar cell populations; the proportion of the anchor cells present was critical (**Fig. 4a/d-g**). Mereu's study used mixed cells of three different species, but focused primarily on the analytical platform, with limited investigation on bioinformatics methods[15]. Nevertheless, as in Tian's study, no non-mixture cells/data were captured. However, studies restricted to mixtures of cell types cannot examine the ability of a method to eliminate variability due to technical factors, important in its capacity to group similar cells together, and also—independently—assess the ability of a method to separate dissimilar cells correctly. We found that analyzing both mixtures of dissimilar cells in various proportions and unmixed samples of the two cell lines provided important additional insights. For example, one widely used method[35] excelled at grouping similar cells together, but when large proportions of two dissimilar cell types were analyzed, this algorithm overcorrected, completely failing to separate B cells from breast cancer cells (**Fig. 4a-g**). Also, as noted previously, it may be difficult to replicate the cellular sample used in Mereu's study for confirmatory or further exploratory evaluations. In contrast, our study was not only empowered by a mixology design, but was also further strengthened by the inclusion of non-mixture sample/data captured separately. Moreover, our reference cell lines are commercially available for future benchmarking studies. Another unique advantage of our standard reference samples is the availability of massive cross-platform deep WGS data (see companion NBT paper), which will be valuable resource for benchmarking future single-cell WGS or proteomics technologies.

In summary, we assessed scRNA-seq data generated with multiple platforms across several centers using samples derived from two well-characterized, biologically distinct cell lines, analyzed either separately or as mixtures. This experimental design allowed us to benchmark the effect of sample cell composition and evaluate variations due to both platform-specific effects and each element of the bioinformatic analysis pipelines, particularly the batch-effect correction algorithms. We believe this will provide a useful resource to the community to benchmark additional scRNA-seq protocols and bioinformatics algorithms. Overall, our study showed that while batch effects were large, the variations across sites and platforms could be corrected by appropriate computational methods. Critically, this study identified the importance of choosing computational methods appropriate to both the technology platforms used and the composition of the samples analyzed. An important conclusion we draw is that the capabilities and limitations of the various elements of the bioinformatic analyses should be chosen to match the experimental situation. A more detailed enumeration of our conclusions is presented in the Online Methods and in the supplementary file (**Suppl. File 1**). The capabilities and limitations of the methods we evaluated are ranked graphically in Figure 6. Best practice recommendations based on our findings are presented in flow chart format (**Suppl. Fig. 25**).

#### METHODS

##### Study design

The schematic overview of the study design is illustrated in **Figure. 1a**. Briefly, two well-characterized reference cell lines (sample A, breast cancer cell line vs. sample B, a matched control B lymphocyte cell line) were used to generate scRNA-seq data across four platforms (10X Genomics, Fluidigm C1, Fluidigm C1 HT, and Takara Bio ICELL8), at four testing sites (LLU , NCI, FDA, and Takara Bio) using standard manufacturer's protocols. At the 10X_LLU and 10X_NCI sites, mixed singe-cell captures and library constructions were also prepared with either 10% or 5% cancer cells spiked into the B lymphocytes, respectively. At the NCI site, single-cell captures and library constructions were also performed with methanol-fixed cell mixtures (5% cancer cell spiked into B lymphocytes, named as fixed_1 and fixed_2 in two sample captures, separately). The 10X scRNA-seq libraries constructed at NCI were sequenced using a shorter modified sequencing method (26x57 bp) at NCI site. One set of 10X scRNA-seq libraries constructed at the NCI site was also sequenced at LLU site using the standard sequencing method (26x98 bp). Bulk cell RNA-seq was also obtained from these cell lines, each in triplicate. All scRNA-seq data were subject to 3 different pre-processing pipelines for either 10X or C1/Takara Bio ICELL8 technologies, respectively. We evaluated eight normalization methods, SCTranform, Scran Deconvolution, CPM, LogCPM, TMM, DESeq, Quantile, and Linnorm and seven batch effect correction algorithms including Seurat v3, fastMNN, Scanorama, BBKNN, Harmony, limma, and ComBat. The cross-platform and cross-center performances were further evaluated by t-SNE, UMAP, and three quantitative metrics (silhouette score, modified alignment score, and kBET), as well as scatter plotting and feature plotting. scRNA-seq data were also compared with population-average RNA-seq data.

**Abbreviations and notations for Fig. 1a: 10X_LLU**, single cells were captured using a 10X Genomics Chromium controller; scRNA-seq was done at the Loma Linda University (LLU) Center for Genomics using the standard 10X Genomics protocol (26x98 bp); **10X_NCI_M**, 10X Genomics scRNA-seq libraries were prepared and sequenced at the NCI sequencing facility using a modified 10X sequencing protocol (26x56 bp); **10X_NCI**, the same 10X Genomics scRNA-seq libraries prepared at the NCI sequencing facility but were sequenced at LLU using the standard 10X sequencing protocol (26x98 bp); **C1_FDA_HT**, single cells were captured using a Fluidigm C1 HT IFC and the scRNA-seq libraries were sequenced at the FDA sequencing facility (75x2 bp); **C1_LLU**, single cells were captured using a Fluidigm C1 IFC chip and the scRNA-seq libraries were sequenced at the LLU Center for Genomics (150x2 bp, ~4-4.77M reads/cell); **ICELL8_PE**, single cells were captured using an ICELL8 chip (Takara Bio) and scRNA-seq libraries were paired end sequenced (75x2 bp) at Takara Bio; **ICELL8_SE**, the same scRNA-seq libraries generated at Takara Bio were also sequenced at the LLU Center for Genomics (150x1 bp, ~1M reads/cell). See Suppl. **Table 1** for details on the numbers of single cells captured and sequencing read depths in each platform and each site.

##### Cell culture and single cell preparation

We obtained the human breast cancer cell line (HCC1395, sample A) and the matched normal B lymphocyte cell line (HCC1395 BL, sample B) from ATCC (American Type Culture Collection, Manassas, VA, USA). The two cell lines were derived from the same human subject (43 years old, female). HCC1395 cells were cultured in RPMI-1640 medium supplemented with 10% fetal bovine serum (FBS). HCC1395BL cells were cultured in IMDM medium supplemented with 20% FBS.

Single cell suspensions were generated by dissociating adherent cells (HCC1395) with Accutase (Innovative Cell Technologies, AT104) or by harvesting suspensions cells (HCC1395 BL). We passed all cells through a 30-micron MACS SmartStrainer (Miltenyi Biotec, 130-098-458) to -remove the cell aggregates.

**Single-cell full-length cDNA generation and RNA-seq using the C1 Fluidigm system**

Single cells were loaded on a medium-sized (10-17 µm) RNA-seq integrated fluidic circuit (IFC) at a concentration of 200 cells/µl. Capture occupancy and live/dead cells at the capture site were recorded using a fluorescence microscope after staining with the live/dead viability/cytotoxicity kit (Life Technologies, L3224). Full-length cDNAs were generated using the Fluidigm C1 system at the LLU Center for Genomics using the SMART-Seq v4 Ultra Low Input RNA kit (Takara Bio) according to the manufacturer's protocol. Only cDNAs generated from live single cells were used for further library construction.

Libraries were prepared using the modified Illumina Nextera XT DNA library preparation protocol. Briefly, the concentrations of cDNAs harvested from the IFC were quantified using the Quant-iT PicoGreen dsDNA Assay (Life Technologies) and then further diluted to 0.1-0.3 ng/µl. 1.25 µl diluted cDNA was incubated with 1.25 µl tagmentation mix and 2.5 µl tagment DNA buffer for 10 minutes (min) at 55 °C. Tagmentation was terminated by adding 1.25 µl of NT buffer and centrifuging at 2,000g for 5 min. Sequencing library amplification was performed using 1.25-µl Nextera XT Index primers (Illumina) and 3.75 µl Nextera PCR Master Mix with 12 PCR cycles. Barcoded libraries were purified and pooled at equal volume. 80 libraries were generated from HCC1395 cells (sample A) and 66 libraries were generated from HCC1395 BL cells (sample B). Library pools were sequenced on the Illumina HiSeq4000 sequencer for 150xbp paired-end sequencing at the LLU Center for Genomics.

**Single-cell 3' End RNA-seq using C1 Fluidigm high-throughput (HT) system**

High-throughput single cell 3' end cDNA libraries were generated according to the manufacturer's instructions at the FDA's Center for Biologics Evaluation and Research. Briefly, single cells were loaded on a HT IFC at a concentration of 400 cells/µl (Nexcelom Cellometer Auto T4). Capture occupancy and live/dead cell at the capture site were recorded using a fluorescence microscope after staining with live/dead viability/cytotoxicity kit (Life Technologies). After cell lysis, the captured mRNA was barcoded during the reverse transcription step with a barcoded primer, and the tagmentation step was done following the Nextera XT DNA library preparation guide. Only polyadenylated RNAs containing the preamplification adapter sequence at both ends were amplified. Lastly, sequencing adapters and Nextera indices were applied during library preparation. Only the 3' end of the transcript was enriched following PCR amplification.

203 libraries were generated from HCC1395 cells (sample A) and 241 libraries were generated from HCC1395 BL cells (sample B). Library pools were sequenced on the Illumina NextSeq 2500, 75xbp, paired-end at the FDA's Genomics Facility.

**Single-cell RNA-seq using the 10X Genomics platform**

After filtering with a 30-micron MACS SmartStrainer (Miltenyi Biotec), single cells were resuspended in PBS (calcium and magnesium free) containing 0.04% weight/volume BSA (400 µg/ml), and further diluted to 300 cells/µl after cell count (Countess II FL, Life Technologies). For the 5% spike-in and 10% spike-in cell mixtures, 5% or 10% of HCC1395 breast cancer cells were mixed with either 95% or 90% of HCC1395BL cells.

Single-cell RNA-seq library preparation was performed following the 3' scRNA-seq 10X Genomics platform protocol using v2 chemistry. Briefly, based on the cell suspension volume calculator table, 3000 cells (17.4 µl of 300 cells/µl suspension) and barcode-beads as well as RT reagents were loaded into the Chromium Controller to generate single Gel Bead-in-Emulsions (GEMs). cDNAs were generated after GEM-RT incubation at 53 °C for 45 min and 85 °C for 5 min. cDNA amplification was performed for 12 PCR cycles following GEM cleanup. After size selection with SPRIselect Reagent, cDNA was incubated for fragmentation, end repair, A-tailing, and adapter ligation. Lastly, sequencing library amplification was performed using sample index primer for 10 cycles.

The methods for other single-cell captures, scRNA-seq library constructions, and sequencing data generation can be found in the **Online Methods** section.

#### ONLINE METHODS

**Methods for other single cell captures, library construction, and sequencing data generation**

**10X Genomics scRNA-seq library construction using fixed cells**

We also constructed 10X scRNA-seq libraries using fixed cells at the NCI site. Briefly, for delayed captures, cells were fixed in methanol using a method described by Alles et al[46]. The fixed samples underwent two different treatments. For the sample of spikein_5%_Fixed_1, the normal and tumor cells were harvested, washed, counted, and a 5% spike-in of breast cancer cells plus 95% normal B cells were prepared and mixed as described above. Approximately 130,000 cells were then processed for fixation. The cells were washed twice with 1X DPBS at 4 °C and resuspended gently in 100µl 1X DPBS (ThermoFisher Scientific). 900 µl chilled methanol (100%) was then added drop by drop to the cells with gentle vortexing. Cells were then fixed on ice for 15 mins, following which they were stored at 4 °C for 6 days. For rehydration, the fixed cells were pelleted by centrifugation at 3000g for 10 mins at 4 °C and washed twice with 1X DPBS containing 1% BSA and 0.4U/µl RNase inhibitor (Sigma Aldrich). The cells were then counted and the concentration was adjusted to be close to 1000 cells/µl. Approximately 8000 cells were loaded onto a single-cell chip for GEM generation using the 10X Genomics

Chromium controller. 3’mRNA-seq gene expression libraries were prepared using the Chromium Single Cell 3′ Library & Gel Bead Kit v2 (10X Genomics) according to the manufacturer guidelines.

For the sample spikein_5%_Fixed_2, breast cancer cells and normal B cells (approximately 4 million each) were harvested and fixed. The cells were initially washed with 1X DPBS and resuspended in 10% 1X DPBS and 90% chilled methanol, as described above. Cells were then fixed on ice for 15 mins, following which they were stored at 4 °C for 24 hrs. For rehydration, the fixed cells were washed with 1X DPBS containing 1% BSA and 0.4U/µl RNase inhibitor and counted. Approximately 8000 cells were loaded onto a single-cell chip for GEM generation using the 10X Genomics Chromium controller. 3’mRNA-seq gene expression libraries were prepared using the Chromium Single Cell 3′ Library & Gel Bead Kit v2 (10X Genomics) according to the manufacturer’s guidelines.

All the 10X Genomics scRNA-seq libraries constructed at the LLU were sequenced on the NextSeq 550 and HiSeq 4000 with the standard sequencing protocol of 26x98 read length at the LLU Center for Genomics, whereas the libraries constructed at the NCI site were either sequenced on the NextSeq 550 with a modified sequencing protocol of 26x57 read length at the NCI Genomics Facility or on the HiSeq4000 using the standard sequencing protocol of 26x98 read length at the LLU Center for Genomics.

**Single-cell RNA-seq using Takara Bio ICELL8 platform**

We also constructed ICELL8 scRNA-seq libraries at Takara Bio USA site on the two cell lines.

ICELL8 Cell preparation and Single Cell Selection

A bulk cell suspension of either cancer or B cells (~ 1 x $10^6$ each) was fluorescently labeled with a premade mix of Hoechst 33324 and Propidium Iodide (Ready Probes Cell Viability Imaging Kit, Thermo Fisher Scientific) in appropriate complete medium for 20 min at 37 °C. Adherent cells were first treated with Accutase as per manufacturer’s instructions (Thermo Fisher Scientific) to dissociate cells from the flask surface. Cells were washed in 1X PBS, (no $Ca^{2+}$, $Mg^{2+}$, Phenol Red, or serum, pH 7.4; (Thermo Fisher Scientific) and centrifuged (100g 3 min) and resuspended in 1 mL of 1X PBS. Cell counts were determined using a Moxie Flow cell counter (ORFLO Technologies, ID, USA) and diluted to ~1 cell in 35 nl (~ 28,600 cells/ml) in 1X PBS (1X PBS, no $Ca^{2+}$, $Mg^{2+}$, Phenol Red, or serum, pH 7.4; Thermo Fisher Scientific) containing Second Diluent (1X), RNase Inhibitor (0.4 U) and 1.92 µM of the 3’ oligo dT terminating primer: SMART-Seq® ICELL8® CDS (Takara Bio USA, CA, USA).

Each cell type solution was dispensed from a 384-well source plate into individually addressable wells in a 5,184 nano-well, 250 nl volume ICELL8 chip (SMARTer™ ICELL8® 250v Chip, Takara Bio USA, CA, USA) using a Multi Sample Nano Dispenser (MSND, SMARTer™ ICELL8® Single-Cell System, Takara). Chip wells were sealed using SmartChip Optical Imaging Film (Takara Bio USA) and centrifuged at 300g for 5 min at 22 °C. All nano-wells in the chip were imaged with a 4X objective using Hoechst and Texas Red excitation and emission filters. Images (TIFF format) were analyzed using automated microscopy image analysis software Cell Select (Takara Bio USA). The chip was stored in a chip holder at -80 °C overnight. Image analysis confirmed cell deposition followed a Poisson distribution. 600 individual nano-wells, each bearing microscopy-identified single live cells, were chosen from each cell type. A well-selection map (filter file) was then autogenerated by Cell Select software to enable individual addressing of the chosen wells for addition of cDNA synthesis and library preparation reagents as detailed in the following sections. All on-chip liquid handling was performed with the MSND. After all dispensing and sealing steps, chips were centrifuged at 3,220g (3 min). All on-chip thermal cycling was performed using a SMARTer™ ICELL8® Thermal Cycler (Takara Bio USA).

In-chip, full-length cDNA synthesis

The ICELL8 chip (containing dispensed samples) was thawed at room temperature for 10 min and centrifuged at 3,220*g* for 3 min at 4 °C. The chip was subsequently incubated at 72 °C (3 min) and immediately placed at 4 °C. RT-PCR mix (35 nl total) was added to each of the previously selected nano-wells (identified as bearing a single cell via the ICELL8 filter file), and the reactions were thermally cycled in-chip as follows: 45.6 °C, 5 sec; 41 °C, 90 min; 99 °C, 9 sec; 95.5 °C, 1 min; 100 °C, 5 sec; 99 °C, 7 sec; (9 °C, 5 sec; 64 °C, 30 sec; 69.5 °C, 5 sec; 67.5 °C, 3 min; GoTo step 5 and repeat 7X, (4 °C hold).

In-chip, P5 index addition and tagmentation

72 primer sequences bearing P5 indices (SMART-Seq® ICELL8® Forward Indexing Primer Set A (5'-AATGATACGGCGACCACCGAGATCTACAC(*i5)*TCGTCGGCAGCGTC-3'); *i5* refers to 1-of-72 unique, 8 nucleotide indices (Hamming distance between P5 indices = 3), were dispensed from a pre-aliquoted 384-well plate in 35 nl aliquots into 72 filter-file identified, nano-well "rows". The chip was sealed with Microseal A film and centrifuged at 3,220*g* (3 min) at 4 °C before returning to the MSND, permitting addition of Tagmentation Master Mix containing: $MgCl_2$, Nextera Amplicon Tagment Mix (Illumina); Terra™ PCR Direct Polymerase Mix, and TRH (Takara Bio USA). The chip was sealed with Microseal A film and recentrifuged as above. Tagmentation was performed in-chip at the following temperatures: 42 °C, 4 sec; 37 °C for 30 min; 4 °C hold.

In-chip, P7 index and PCR reagent addition: first PCR generating 5,184 unique indices

A reagent mix containing 72 primer sequences bearing P7 indices (SMART-Seq® ICELL8® Reverse Indexing Primer Set A, 5'-CAAGCAGAAGACGGCATACGAGAT(*i7)*GTCTCGTGGG CTCGG-3'); *i7* refers to 1-of-72 unique, 8 nucleotide indices (Hamming distance between P7 indices = 3), were dispensed from the same pre-aliquoted 384-well index plate (separate location for P7 indices) in 35 nL aliquots, into 72 filter file identified "columns" of the chip. As a consequence of adding separate P5 and P7 indices to rows or columns, a 72 x 72 *m x n* matrix of combinatorial P5 and P7 pairs was generated, uniquely identifying each of the 5,184 nanowells. The chip was sealed with SmartChip Sealing Film and centrifuged at 3,220*g* for 3 minutes at 4 °C. PCR cycling was performed as follows: (77 °C, 12 sec); (72 °C, 3 min); (99 °C, 11 sec); (95.5 °C, 1 min); (100 °C, 20 sec); (99 °C, 10 sec); (53.3 °C, 5 sec); (58 °C, 15 sec); (71 °C, 5 sec); (67.5 °C, 2 min; (Go To step5 and repeat 7X); (4 °C, hold).

Off-chip, sample extraction and purification of round 1 PCR amplicons

Round 1 PCR amplicons were collected from the ICELL8 chip using the SMARTer ICELL8 Collection Kit: Collection Fixture, Collection Tube, and Collection Film into a collection and storage tube as per manufacturer's instructions (Takara Bio USA). 50% of the extracted library was purified twice using a 1X proportion of AMPure XP beads (Beckman Coulter) to a final volume of 14 µl in Elution Buffer, provided with the SMART-Seq ICELL8 Reagent Kit.

Off-chip, library amplification (2nd PCR)
Double-AMPure bead-purified, first round amplicon (14 µl, from above) was PCR amplified in a 50 µl volume of 2$^{nd}$ PCR Mixture containing SeqAmp™ CB PCR Buffer (25 µl), 5X Primer Mix (P5 and P7 primers) and Terra™ PCR Direct Polymerase Mix 0.05 U/ µl final concentration (Takara Bio USA) via a thermal protocol: (98 °C, 2 min) x1; followed by 8 thermal cycles: 98 °C, 10 sec; 60 °C, 15 sec; 68 °C, 2 min. This sequencing-ready NGS library was purified using 1 round of a 1X proportion of AMPure XP beads (Beckman Coulter). The final elution volume was 17 µl in Elution Buffer.

ICELL 8 scRNA-seq library QC and Sequencing
The NGS library concentration (ng/µl) was determined using a Qubit fluorometer (Thermo Fisher). Based on Qubit readings, 1-2 ng/µl was examined using a 2100 Bioanalyzer and a corresponding High Sensitivity DNA Kit (Agilent) to determine the MW profile of the size-selected library. The Bioanalyzer amplicon sizes ranged between 200 to 3000 bp, with an average size of 550 bp. The ICELL scRNA-seq libraries were sequenced both at the Takara Bio USA site on Illumina NextSeq 550, 75xbp, paired-end and at the LLU Center for Genomics on HiSeq 4000, 150xbp, single-end.

**Bulk cell RNA-seq**

We isolated bulk-cell total RNA from the HCC1395 and HCC1395 B cells using miRNeasy Mini kit (QIAGEN), and constructed RNA-seq libraries using the NuGEN Ovation universal RNA-seq kit at LLU. Briefly, 100 ng of total RNA was reverse transcribed and then converted into double stranded cDNA (ds-cDNA) by addition of a DNA polymerase. The ds-cDNA was fragmented to ~200 bps using the Covaris S220, and then underwent end repair to blunt the ends followed by barcoded adapter ligation. The remainder of the library preparation followed the manufacturer's protocol. All the libraries were quantified using Qubit 3.0 (Life Technologies) quality checked on a TapeStation 2200 (Agilent Technologies). The bulk-cell RNA-seq libraries were sequenced both on a NextSeq 550, 75xbp paired-end; and on a HiSeq 4000, 100xbp paired-end at the LLU Center for Genomics.

**Overall data generated, and data QC assessments**

**Supplementary Table 1** summarizes the overall cell numbers and sequencing reads of single cells captured across all four sites, a total of twenty different scRNA-seq datasets. **31,887** single cells with either 3' or full-length scRNA-seq data were captured (**Suppl. Table 1 & Suppl. Fig. 1**). Five libraries from the NCI site (10x_NCI_M) were also re-sequenced using a modified sequencing protocol (26x57 bp). Across all the platforms and datasets, over 93.6% of the reads were mapped to the exonic and non-exonic regions except for sample A of 10X_NCI_M (modified shorter sequencing), which had a mapping rate of 87% (sample A) and 90.3% (sample B) (**Fig. 1b**). However, there were variations in the mapping rates to exonic regions across platforms and sites, with ICELL8 and Fluidigm C1 full-length transcript methods showing a higher mapping rate than 3'-transcript scRNA data in the case of tumor cells (sample A: C1_LLU_A, 83.1%; ICELL8_PE_A, 84.0%; ICELL8_SE_A, 80.7% vs. 10X_LLU_A, 65.3%;10X_NCI_A; 66.3% for the 10X data). The UMI (unique molecular identifier) data generated by the 10X platform showed that 35.3% of the mapped reads (or 57.0% of the exonic reads) were derived from de-duplicated UMIs in breast cancer cells (sample A), and 22.2% of the mapped reads (or 35.2% of the exonic reads) were derived from de-duplicated UMIs in normal B cells (sample B). We also noticed that the exonic mapping rates were slightly lower for the 10X Genomics technologies when using the modified shorter sequencing protocol compared with the standard sequencing protocol (26x57 bp vs. 26x98 bp). The modified protocol used a shorter sequencing read length, which could cause a higher percentage of non-specific mapping reads. Nevertheless, many overlapping genes were detected (96.6%-97.3%) with a high correlation (R=0.997-0.998) between the standard and modified sequencing protocol for the 10X Genomics scRNA-seq (**Suppl. Fig. 2**).

**Bioinformatics methods**

**Reference genome:** The reference genome and transcriptome were downloaded from the 10X website as refdata-cellranger-GRCh38-1.2.0.tar.gz, which corresponds to the GRCh38 genome and Ensmebl v84 transcriptome. All the following bioinformatics data analyses are based on the above reference genome and transcriptome.

**Preprocessing of UMI based scRNA-seq data from the 10X platform**

For UMI based 10X samples, three pre-processing pipelines, CellRanger (v3.1.0), umitools[21] (v1.0.0), and zUMIs[22] (v2.4.5) were used to process the raw fastq data and generate gene count matrices. In the CellRanger pipeline, cellranger count was used with all default parameter settings. In the umitools and zUMIs pipelines, reads were filtered out if phred sequence quality of cell barcode bases was < 10 or UMI bases < 10. In the zUMIs pipeline, option -d was used to perform downsampling analyses to 8 fixed depths (5k, 10k, 25k, 50k, 100k, 150k, 200k, and 250k) to generate gene count tables. With umitools, umi_tools whitelist with default parameter settings was used to generate a list of cell barcodes for downstream analysis. umi_tools extract was used to extract the cell barcodes and filter the reads (options: --quality-filter-threshold=10 --filter-cell-barcode). STAR (v2.5.4b)[28] was used for alignment to generate bam files containing the unique mapped reads (option: outFilterMultimapNmax 1) for gene counting. featureCounts (v1.6.1)[23] was used to assign reads to genes and generate a BAM file (option: -R BAM). samtools (v1.3)[47] sort and samtools index were used to generate sorted and indexed BAM files. Finally, umitools count (options: --per-gene --gene-tag=XT --per-cell --wide-format-cell-counts) was used for the *sorted BAM files to generate gene count per cell matrices*

**Preprocessing of non-UMI based scRNA-seq data from C1 and Takara Bio ICELL8 platforms**

For non-UMI based samples, three pre-processing pipelines were compared for processing the raw fastq data and generating gene count matrices. The pipelines included trimming and filtering, alignment, and gene counting. In the trimming and filtering process, one of the three tools [Trimmomatic (v0.35)[27], trim_galore (v0.4.1)[48], or cutadapt (v1.9.1)[26]] was used to process the raw fastq data. Bases with quality less than 10 were trimmed from 5' and 3' ends of reads. Reads less than 20 bases were excluded from further analysis. STAR with default parameter settings was used for alignment to generate bam files. Three gene counting tools, featureCounts, RSEM (v1.3.0)[25], or kallisto (v0.43.1)[24] were used to generate gene counts per cell. All default parameter settings were used except the following: In RSEM, option --single-cell-prior was used to estimate gene expression levels for scRNA-seq data; Option of --paired-end was used if the data were paired-end fastqs; In kallisto, options -l 500 and -s 120 were used to represent estimated average fragment length and standard deviation of fragment length if the data were single-end fastqs.

**Preprocessing and differential gene expression analysis of bulk RNA-seq data**

The preprocessing pipeline of bulk RNA-seq data included QC (FastQC v0.11.4), trimming and filtering (Trimmomatic), alignment (STAR), and gene counting (RSEM). The parameter settings in the pipeline were the same as the preprocessing pipelines used for non-UMI scRNA-seq data. In RSEM, the option --single-cell-prior was turned off for estimation of gene expression levels in bulk RNA-seq data. DESeq2 (v1.24.0) was used to perform the differential expression analysis between breast cancer samples and normal samples with default parameters.

**BGL and data sharing within the team**

Working under the FDA single-cell sequencing consortium, to streamline fast pre-processed data sharing, access, and analysis, we used the BioGenLink™ (BGL) platform from Digicon Corporation as a central repository to host the pre-processed data as described above. All data including the single-cell RNA-seq data were pre-processed at LLU and then the pre-processed data were either uploaded into BGL from LLU Center for Genomics servers or using tools within BGL that utilized Globus, file transfer protocol (FTP), and secure copy protocol (SCP). Detailed data annotation files about all genomics data were also uploaded into the BGL.

**Performance of normalization methods across all datasets**

We investigated some existing bulk RNA-seq normalization procedures including "Counts per Million (CPM)", "Trimmed Mean of M values (TMM)", "Upper Quantiles", "DESeq" normalization implemented in the DESeq Bioconductor package, and "Trimmed Mean of M values (TMM)" implemented in edgeR. There were also methods that were specifically tailored to scRNA-seq data sets, such as SCTransform, scran, and Linnorm. Both scran and Linnorm were run using default parameters. SCTransform was run without regressing out any variables with default setting.

We performed reads downsampling of each cell to two different read depths (10K and 100K per cell) for each data set and evaluated the performance of the normalization methods of two read depths per data set. Similar to the method used in the *scone* paper[49], the metric we used to assess normalization methods was based on how well the two samples from the same cell were grouped with each other. In detail, we used silhouette width, which is defined as,

$$s(i) = \frac{b(i) - a(i)}{max\,\{a(i), b(i)\}}$$

For each cell *i*, let *a(i)* be the average distance between *I* and all other cells within the same cluster. Let *b(i)* be the lowest average distance of *I* to all points in any other cluster, of which *I* is not a member. Here we defined the clustering structure that the same cells from two different sequencing runs form a single cluster, thus we have a total number of n/2 clusters if the total number of samples is n.

We calculated the silhouette width values of each dataset. The larger the silhouette width values, the better the performance of the normalization method.

**scRNA-seq data batch effects and batch-effect correction pipelines**

We used the gene count matrix from the Cell Ranger pipeline (10X Genomics data) and STAR-featureCounts pipeline (non 10X Genomics data) as input to evaluate batch correction methods. In the Cell Ranger pipeline, both CellRanger 2.0 and CellRanger 3.1 were applied to 10X Genomics data. The batch correction evaluation of the data processed by CellRanger 2.0 was provided as supplementary figures. For the evaluation, three different conditions were considered as (1) all data sets; (2) data sets with biologically similar cells; and (3) data sets with biologically different cells. The evaluation procedure included the following four major steps:

1. Monocle2[50, 51] strategy to filter dead cells and doublets for 10X Genomics single cell data
2. Single-cell data processing and highly variable gene (HVG) selection
3. Batch correction by seven different methods
4. Evaluation by *t*-SNE or UMAP, kBET (kBET v0.99.5) acceptance score, modified alignment score, and silhouette score.

A detailed description and functions used for batch correction are summarized in **Supplementary Table 9**.

In step 1, all 10X single cell data sets were processed by the Monocle2 strategy to filter dead cells and doublets. In this strategy, the total numbers of UMIs and genes for each cell were counted. The upper bound was calculated as mean plus two standard deviations (SD) and the lower bound as mean minus two SD for both the total UMIs and genes, respectively. Cells with total UMIs or genes outside of the upper and lower bounds were removed.

In step 2, Seurat (v3.0.3) based data processing was applied to each data set. Genes detected in fewer than 3 cells and cells containing less than 200 genes were removed from the data sets prior to further analysis. The data sets were then log transformed and scaled. The top 2,000 HVGs were selected in each data set with function *FindVariableGenes* for the five R-based batch correction methods Seurat, fastMNN (scran v1.12.1 and

SeuratWrappers v0.1.0), Harmony (v0.99.9), limma (v3.40.4), and ComBat (v3.32.1). For the Python-based batch correction methods Scanorama (v1.4) and BBKNN (v1.3.5), the detailed description of data processing is provided in the 'Scanorama processing' and 'BBKNN processing' sections.

In step 3, the processed data and HVGs in step 2 were used as input to perform batch correction. The main functions and parameter settings of the seven batch correction methods are summarized in **Supplementary Table 9.**

In step 4, the t-SNE and UMAP plots and the calculations of the kBET acceptance score, modified alignment scores, and silhouette scores were based on the low-dimensional embedding matrices of each batch correction method. For Seurat v3, fastMNN, and Harmony, Seurat and SeuratWrappers were used to generate low-dimensional embedding matrices. Seurat based PCA reduction was applied to batch corrected matrices by Scanorama, limma, and ComBat to generate low-dimensional matrices, whereas for BBKNN, the UMAP coordinates matrices were used as the low-dimensional embedding matrices. The functions used to generate tSNE and UMAP can be found in **Supplementary Table 9.**

**Scanorama pipeline**

The Scanorama Python package was used to process the data sets and perform batch correction. The script *process.py* with default parameters was used to perform cell filtering and normalization. 2000 HVGs were used in the function *correct* to perform batch correction and generate Scanorama-corrected gene expression matrices.

**BBKNN pipeline**

The Seurat-inspired Scanpy (v1.4.4) Python workflow was applied to process the data sets. All data sets were input using the function *pd.read_csv* in the pandas package, transferred into annotated data matrices, and appended into a list using the function *anndata.AnnData* from the package anndata. Cells and genes were filtered using functions *scanpy.api.pp.filter_cells* and *scanpy.api.pp.filter_genes* with the same parameter settings as at Step 1. The processed data matrices were merged to generate a master gene expression matrix and further log transformed and normalized by functions *scanpy.api.pp.log1p* and *scanpy.api.pp.normalize_per_cell*. The top 2,000 HVGs were selected from the merged gene expression matrix by the function *filter_genes_dispersion* with the same parameter settings as at Step 2. Further log transformation (function *scanpy.api.pp.log1p*) and scaling (function *scanpy.api.pp.scale*) were performed for the newly generated gene expression matrices. The function *bbknn* with default parameters was carried out for the batch correction.

**FastMNN vs. MNN**

In **supplementary figure 16**, we further compare the performance of fastMNN and MNN. The steps to perform MNN correction are the same as fastMNN except the batch correction (step 3). We used the function *mnnCorrect* (scran package v1.8.4) with default parameters to perform the batch correction.

**Preprocessing and batch-effect correction on Tian et al. data[16]**

We preprocessed Tian's data with the CellRanger pipeline for 10X data and umitools pipeline for their non-10X data. The same procedures for the seven batch correction methods described previously were applied to the preprocessed data to perform batch correction evaluation.

**Bioinformatics pipelines validated and performed in BGL (Biogenlink)**

We carried out some bioinformatics pipelines in BGL to cross-validate some of our bioinformatics data analyses. Bioinformatics tools were created in BGL for performing batch correction of single-cell RNA-seq data using Seurat v3, fastMNN, Scanorama, BBKNN, Harmony, limma, and Combat procedures and for visualizing the results of each procedure using t-SNE and UMAP for scenario # 1. For each procedure, a tool was created in BGL that allows a user to point and click to select input data and parameters for running methods from one or more packages. For each tool, BGL ran a script on the back-end to execute the steps described below. Unless otherwise stated, all functions and procedures used default settings.

**Silhouette width to quantify batch-effect correction**

Silhouette width score of each cell was calculated based on the two cell types, HCC1395 and HCC1395BL, for the scenarios #1 and #4 (**Fig. 4a/d**) by the function *silhouette* from the R package cluster (v.2.0.8). We further calculated the average silhouette width scores of the cells in each cluster, respectively. Finally, the mean of the average silhouette width score was used to represent the performance of the seven batch correction methods.

**kBET acceptance score to quantify batch-effect correction**

kBET acceptance score was calculated using Buttner et al. pipeline[42] for four different sample combination scenarios (**Fig. 4a-d**) to assess the batch correction performance. This metric was calculated based on the low-dimensional embedding matrices of each batch correction method. For Seurat v3, fastMNN, and Harmony, both Seurat and SeuratWrappers were used to generate low-dimensional embedding matrices. Seurat-based PCA reduction was applied to batch corrected matrices by Scanorama, limma, and ComBat to generate low-dimensional matrices, whereas for BBKNN and UMAP, coordinate matrices were used as low-dimensional embedding matrices for the evaluation. The score was calculated for either breast cancer cells or B lymphocytes across different batches, respectively.

**Modified alignment score to quantify batch-effect correction**

We adopted the idea of alignment score from the Butler et al. method[7] to calculate alignment score based on the cells' embedding in two-dimensional space constructed by t-SNE or UMAP. Like kBET, this metric was also calculated based on the low-dimensional embedding matrices of each batch correction method as described in kBET. The score was calculated for either breast cancer cells or B lymphocytes across different batches of scRNA-seq datasets, respectively, for each of four sample combination scenarios (**Fig. 4a-d**). However, due to the difference in cell numbers across different data sets in our study, we developed a modified alignment score calculation algorithm as follows:

1. Calculate the percentage of cells in each data set $i$ as $w_i$ ($i = 1 \dots N$, $N$ is the total number of data sets).
2. For each cell $j(j = 1 \dots N_j)$ of data set $i$, calculate how many of its $k$ nearest-neighbors belong to the same data set as $x_{ij}$ and then take an average of $x_{ij}$ in data set $i$ to get $\acute{x}_i$.
3. Alignment score = $\sum_{i=1}^{N} w_i \left(1 - \frac{\acute{x}_i - w_i k}{k - w_i k}\right)$
4. We chose $k$ to be 1% of the total number of cells, as recommended by Butler et al[7].

**Bioinformatics evaluation of global and cell-type specific gene expression consistency across platforms/sites using all scRNA-seq data**

To investigate the consistency of global gene expression across different platforms/sites and scRNA-seq data sets, we selected benchmarking genes according to the average gene expression (log2(TPM+1)) of bulk RNA-seq (three biological replicates) from samples A and B. We excluded the top 0.1% highly expressed genes to avoid abnormally expressed genes. To obtain the robust genes, we further filtered out genes with standard deviation of gene expression greater than 1 across three replicates to obtain the robust genes. The remaining genes were used to define three different expression groups by selecting the top 500 most highly expressed, 500 intermediately expressed, and 500 rarely expressed genes based on the ranking of average gene expression levels. For the 1500 genes selected, we calculated cell percentage per gene by defining the percentage of cells with the expressed gene (gene counts >= 1) for different scRNA-seq data sets. To get comparable cell percentages, we considered only gene count matrices from the downsampling results (100K reads per cell) of zUMIs (10X data sets) and featureCounts (non-10X data sets) pipelines. The Pearson correlations of the cell percentages between any two scRNA-seq platforms were calculated for each of the three expression groups to evaluate the consistency.

**Scatter plotting**

To assess the variation of gene expression across different platforms, we generated scatterplot matrices. For all platform-specific data sets, which include 7 single cell datasets and 3 bulk cell RNA-seq datasets for each cell line, the raw gene count matrices were converted to normalized gene lists $L^{(i)}$ by computing the average gene expression count $G_{mj}^{(i)}$ of all cells $N$.

$$L_m^{(i)} = \frac{1}{N}\sum_{j=1}^{N} G_{mj}^{(i)}, \ (m = 1, \dots, M) \ \text{Where } G_{mj}^{(i)} = log\left(CPM_{mj}^{(i)} + 1\right).$$

$$L^{(i)} = \begin{pmatrix} L_1^{(i)} \\ \vdots \\ L_M^{(i)} \end{pmatrix}, \qquad (i = 1, \dots, 8)$$

For $i = 1, \dots, 8$ of gene list $L^{(i)}$ , this gives 8 columns which can be grouped to an $M$ x 8 matrix as

$$A = \begin{pmatrix} L_1^{(1)} & \dots & L_1^{(8)} \\ \vdots & \vdots & \vdots \\ L_M^{(1)} & \dots & L_M^{(8)} \end{pmatrix}$$

The final normalized matrix A was used as input to generate scatterplot figures by the R packages ggplot2 and psych. Scatterplot displays read count distributions across all genes and all protocols. Each gene is represented as a point in each scatterplot; $x, y$ values represent the gene expression variation in a pair of protocols compared. In addition, each sample's gene expression distributions are computed and displayed in a bar chart. Pearson correlation coefficient between any pair of the protocols were calculated to show the consistency.

**Violin plotting**

To assess the scRNA-seq gene expression profiles across different platforms based on 4 different RNA groups including protein coding RNAs, antisense RNAs, lincRNAs, and miscRNAs, we took the raw gene count matrices for each data set and converted them to a normalized gene list $L^{(i)}$ by computing the average gene expression count $G_{mj}^{(i)}$ of all cells. Please see scatter plotting normalized count matrix computing method for details. The genes which had expression of zero were removed from the comparison; the filtered gene expression lists were used to extract the specific RNA group genes to generate violin plots using R version 3.6.0, ggplot2_3.2.0 and dplyr_0.8.3 packages.

**Feature plotting**

For the uncorrected data, Seurat objects from t-SNE dimensional reduction were used as the data source for generating feature plots. A total of 20 genes (10 for Sample A, cancer cells; 10 for Sample B, B cells) were selected as cell-type specific markers for Sample A and B based on both the bulk-cell RNA-seq DEG ranking and literature. Each cell was assigned a "CellType" (A if the cell came from cancer cells, B if the cell came from B cells). The Seurat function *FeaturePlot* with default parameters was run to generate the gene expression feature plots, in which each cell was colored based on the expression level of the selected gene.

**Bioinformatics methods for single-cell detection consistency of cell-type specific markers CD40, CD74, and TPM1**

To examine the consistency of three marker genes across different single cell platforms, we used the normalized gene expression data (CPM value) from the downsampling results (100K reads per cell) of zUMIs (10X data sets) and featureCounts (non-10X data sets) pipelines. The expression matrix of three marker genes per cell was generated. The cell percentages with detectable, low, intermediate, or high expression were defined by the percentage of cells with CPM > 0, 0 < CPM < 1, 1 ≤ CPM < 10, and CPM ≥10.

**Methods used to generate summary benchmarking statistics for the various bioinformatics pipelines**

The performance of the various pipelines regarding preprocessing, normalization and batch-effect correction is summarized in **Figure 6** based on a Z-score statistic calculated for each metric as detailed below. To benchmark preprocessing methods in terms of gene detection, we first grouped the fourteen pairwise datasets representing either the normal B cell line or the breast cancer cell line (**Fig. 1b**) into three categories: those processed using the 10X protocol (6 datasets), 3' end counting using the Fluidigm high-throughput protocol (2 datasets), and full-length-based protocols (6 datasets). For each dataset, we calculated the proportion of the number of genes detected per pipeline compared with the maximum number of genes detected for that group. Then, for each pipeline, the average scaled ratios of the detected genes within the three categories were calculated. Finally, a Z-score was calculated based on the average scaled ratios per category per preprocessing pipeline. To assess clusterability following normalization, we determined Z-scores for both the median and variance of the calculated silhouette width scores of the fourteen paired B cell and tumor cell datasets as depicted in (**Fig. 3**). Batch-effect correction performance was assessed in terms of clusterability (ability to separate different cell types from each other) and mixability (ability to group similar cells together across datasets). To assess clusterability following batch-effect correction, a Z-score was derived from the harmonic means calculated for the silhouette width scores obtained from the datasets combining both scenario #1 (**Fig. 4a**, combination of all datasets in a single analysis) and scenario #4 (**Fig. 4d**, spiked-in data). To assess mixability following batch-effect correction, a Z-score was derived from the harmonic means of the kBET acceptance scores from datasets taken across all four tested scenarios (**Fig. 4a-d/g**).

**Conclusions and best practice recommendations**

**1)** There were large variations across different scRNA-seq platforms and centers. **2)** Different pre-processing methods/pipelines detected different numbers of genes and/or cells. **3)** Normalization algorithms alone could not remove the batch effects. **4**) Different normalization strategies performed differently across datasets, protocols and platforms; most performed well for either 3'- or full-length-transcript scRNA-seq platforms such as SCTransform, scran, logCPM, and Linnorm, but TMM and Quantile performed poorly, and are not recommended. **5)** Seurat v3, Harmony, BBKNN, fastMNN, and Scanorama all could correct and remove the batch variations in specific sample and dataset scenarios; we recommend users apply appropriate batch-effect correction methods depending on the characteristics of their datasets (e.g., cellular/sample heterogeneity and composition, platforms used, see **Supplementary Fig. 25**). **6**) BBKNN, fastMNN, and Harmony ranked best for clusterability/cell type identification, whereas Seurat v3, Harmony, and fastMNN performed best for mixability. **7)** fastMNN, BBKNN, and Harmony remove batch variations well across different platforms, including both mixed and non-mixed distinct samples, but the order of importing the datasets into the pipeline and the requirement for a mixed sample are critical for MNN/fastMNN; whereas BBKNN and Harmony worked well regardless of the inclusion of mixed heterogeneous biological distinct samples across platforms and batches; thus, for MNN/fastMMN, we recommend including a mixed sample and importing the mixed data first into the pipeline. **8)** CCA/Seurat v3, despite its superior mixability for biologically similar samples, will over-correct batch effects and misclassify cells (i.e., poor clusterability/cell type identification) if large proportions of distinct cell types are co-present. However, Seurat v3 works well both for clusterability and mixability for datasets when only a small fraction of dissimilar cells (e.g., 5-10%) is present. Thus, we do not recommend using CCA/Seurat v3 for scenarios containing large fractions of biologically distinct cell type samples. **9)** BBKNN performed best in clusterability/cell type identification, but it performs poorly in mixability particularly in heterogeneous cellular samples. **10)** The current version of Scanorama worked well for the 10X Genomics data only, but did not work for non-10X platforms, thus we do not recommend it for non-10X data. **11)** We observed good consistency between CellRanger 3.1 and 2.0 pre-processed data, however, CellRanger 3.1 can detect some extra cells with very few transcripts called, which may affect batch-effect corrections in certain scenarios.

**Conflict of interests and disclaimer:** Andrew Farmer and Alain Mir are employees of Takara Bio USA, Inc., and Ben Ernest and Urvashi Mehra were employees of Digicon Corporation. All other authors claim no conflicts of interest. The views presented in this article do not necessarily reflect current or future opinion or policy of the US Food and Drug Administration. Any mention of commercial products is for clarification and not intended as an endorsement.

**Authors' contributions:** CW and WX conceived and designed the study. CW managed the project and directed bioinformatics data analyses. CW drafted manuscript and annotated all the results. MMJ helped edit the manuscript. WC, CW, BT, MM, MMJ, AF, and AM performed single-cell culturing, single cell captures, scRNA-seq libraries and sequencing. XC, ZWY, YMZ, XJX, VC, YTB, BE, WX, UM, JL, JLL, and CW performed bioinformatics data analyses. WC, XC, ZWY, YMZ, YTB, XJX, VC, MM, AM, MMJ, and JJL prepared the methods for the manuscript. ZWY drew all the figures, WC and HC prepared the tables. CW, MMJ, WC, AF, WX, and YMZ revised the manuscript. All authors reviewed the manuscript. CW finalized and submitted the manuscript.

**Acknowledgements:** The authors would like to thank Ms. Diana Ho of the LLU Center for Genomics for her great administrative support, particularly in coordinating the weekly Zoom conference calls and assistance in preparation of meeting minutes for the FDA SEQC-2 single-cell sequencing project. The authors would like to thank Dr. Wendell Jones at $Q^2$ Solutions | EA Genomics for critical review and helpful comments. The authors would like to thank Dr. Zhong Chen at LLU and Jyoti Shetty at NCI for technical assistance in performing sequencing; John Bettridge at NCI for technical assistance in 10X Genomics scRNA-seq library preparation; Vyacheslav Furtak at FDA for library preparation; Wells Wu at the FDA/CBER Core Facility for Illumina sequencing. The authors also would like to thank Sangeetha Anandakrishnan of Takara Bio USA, Inc. for technical assistance with Takara Bio ICELL8 single cell capture and library preparation. The genomic work carried out at the LLU Center for Genomics was funded in part by the National Institutes of Health (NIH) grant S10OD019960 (CW), the Ardmore Institute of Health grant 2150141 (CW) and Dr. Charles A. Sims' gift to LLU Center for Genomics.

**Software and code availability statement**

We used many algorithms and code for batch correction which have been published previously. All of our code is provided in GitHub and Code Ocean at the following links. We would like to note that Code Ocean link is still under validation and we will provide an update immediately when available.

https://github.com/oxwang/fda_scRNA-seq

https://codeocean.com/capsule/0497386

#### Data availability statement

The datasets generated during and/or analyzed during the current study are available in the SRA repository with the access code # (Sub4635070) and the data can be accessed when the paper is published. The following is the reviewer link which only contains metadata information per the SRA policy:

https://dataview.ncbi.nlm.nih.gov/object/PRJNA504037?reviewer=mv5tvl7jnfaceln7lv354mchg2
