## Supplementary File 2 for "A Multi-center Cross-platform Single-cell RNA Sequencing Reference Dataset"

**Towards Best Practice in Single-cell Sequencing:**

**A Comprehensive Multi-Center Cross-platform Benchmarking Study of Single-cell RNA Sequencing Using Reference Samples**

Wanqiu Chen[1#], Yongmei Zhao[2,8#], Xin Chen[1,3#], Zhaowei Yang[4,1#], Xiaojiang Xu[5], Yingtao Bi[6], Vicky Chen[2,8], Jing Li[4], Hannah Choi[1], Ben Ernest[7], Bao Tran[8], Monika Mehta[8], Parimal Kumar[8], Andrew Farmer[9], Alain Mir[9], Urvashi Mehra[7], Jian-Liang Li[4], Malcolm Moos Jr.[10], Wenming Xiao[11]*, Charles Wang[3,1]*

### Supplementary Figures

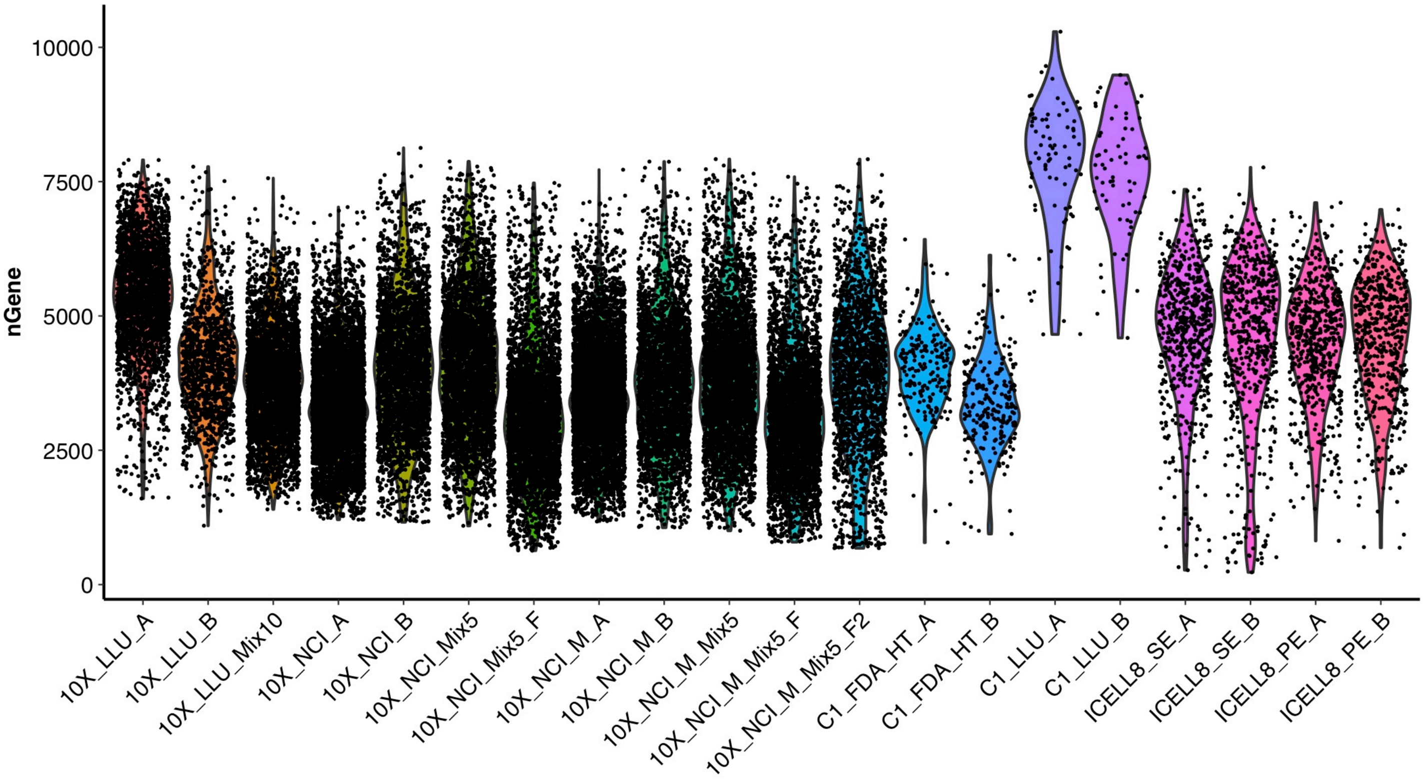


**Supplementary Figure 1. An overview of the number of genes detected in each cell across all platforms/datasets.**

The violin plot shows the number of genes detected in each cell across twenty scRNA-seq datasets. The plot was generated using Seurat (version 3.1). Each dot represents the number of the detected genes for an individual cell. The violin shapes with colors summarize data distributions. X-axis represents scRNA-seq dataset, and Y-axis shows the number of genes that were detected in a cell. The average number of genes detected in each cell is about 4000 and most of the cells had roughly around 2500-7500 genes, except for samples C1_LLU_A and C1_LLU_B. The 10X Genomics scRNA datasets were preprocessed using CellRanger 3.1.

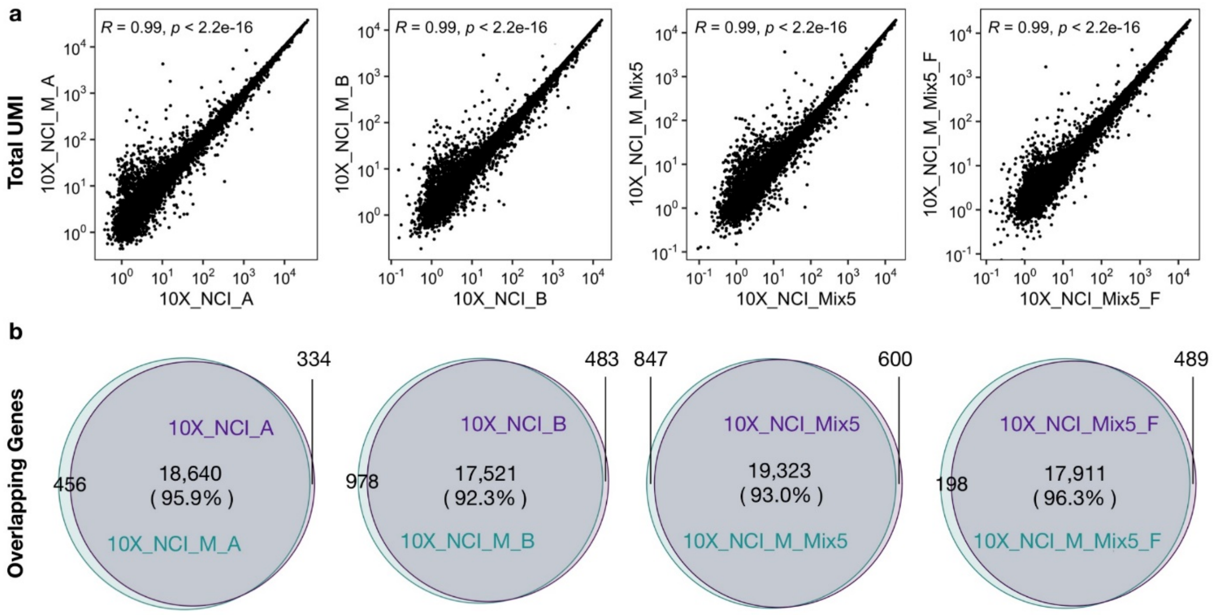


**Supplementary Figure 2. Correlation and overlapping genes between standard and modified sequencing protocols for the 10X Genomics scRNA-seq datasets.**

**(a)** Scatter plots showing the correlation between the standard and modified sequencing protocols for the 10X Genomics scRNA-seq datasets. The total number of unique molecular identifiers was calculated for each gene that occurred in both datasets. The coefficient of correlation (R) of each paired dataset is shown on the top of the scatter plots. **(b)** Venn diagram showing the overlapping genes detected using the standard (26x98 bp) and modified (26x57 bp) sequencing protocols. Four paired datasets were used for the comparison: 10X_NCI_A and 10X_NCI_M_A captured from breast cancer cells (HCC1395), 10X_NCI_B and 10X_NCI_M_B captured from B cells (HCC1395BL), 10X_NCI_Mix5 and 10X_NCI_M_Mix5 captured from B cells with 5% cancer cells spiked-in, and 10X_NCI_Mix5_F and 10X_NCI_M_Mix5_F captured from B cells with 5% cancer cells spiked-in, which had both been methanol fixed. CellRanger 3.1 was used to pre-process the datasets.

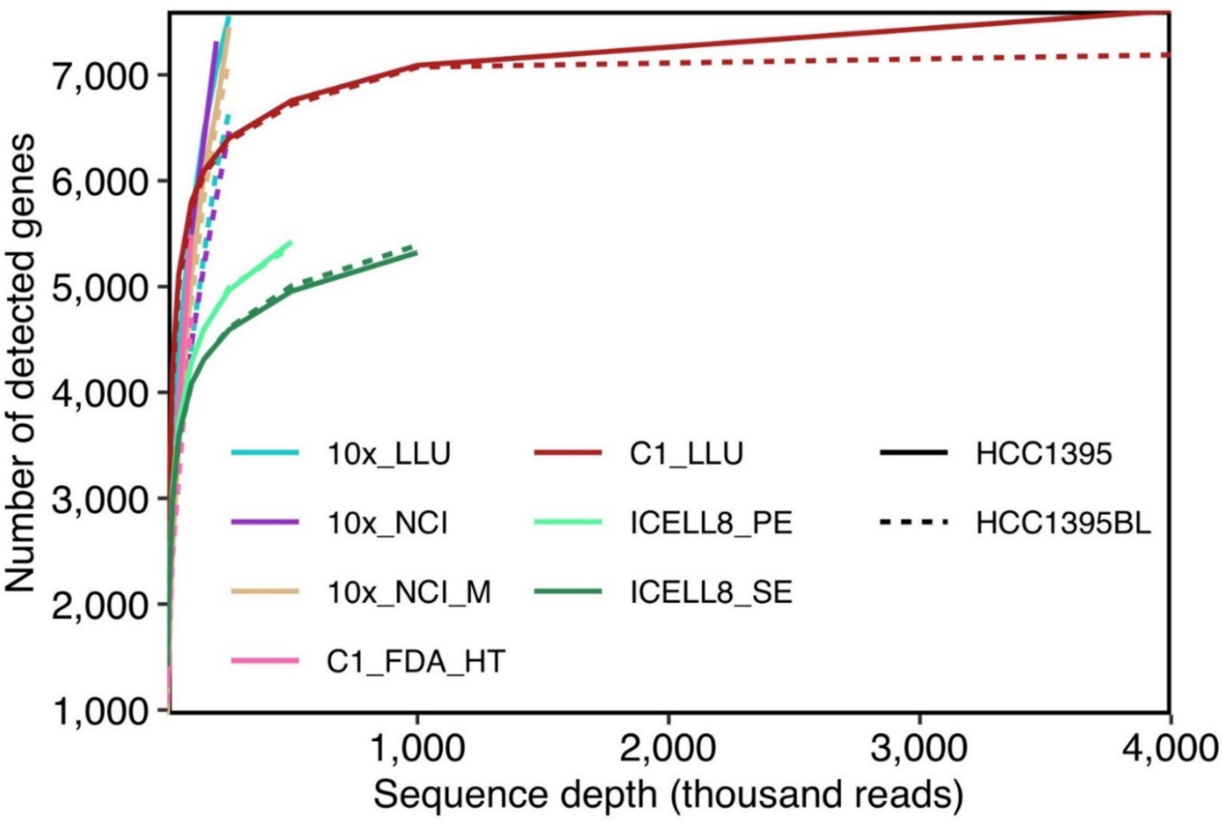


**Supplementary Figure 3. Median number of genes detected per cell at different sequencing read depth across scRNA-seq datasets.**

Fourteen datasets, including seven scRNA-seq data from breast cancer cells (HCC1395) and seven scRNA-seq from B cells (HCC1395BL), were evaluated. Because of the low cell number captured by the C1 platform, over 4000 K reads per cell were generated**.** For all the 10X- and C1_FDA_HT- derived 3' scRNA-seq datasets, the sequence depth was ~100 K reads per cell due to the higher number of cells analyzed. For the ICELL8 platform, we generated ~ 200 K reads per cell for paired-end sequencing and ~ 1000 K reads per cell for single-end sequencing. The solid line represents the breast cancer cells (HCC1395); the dashed line represents the B lymphocytes (HCC1395BL). All the 10X data were preprocessed by CellRanger 3.1.

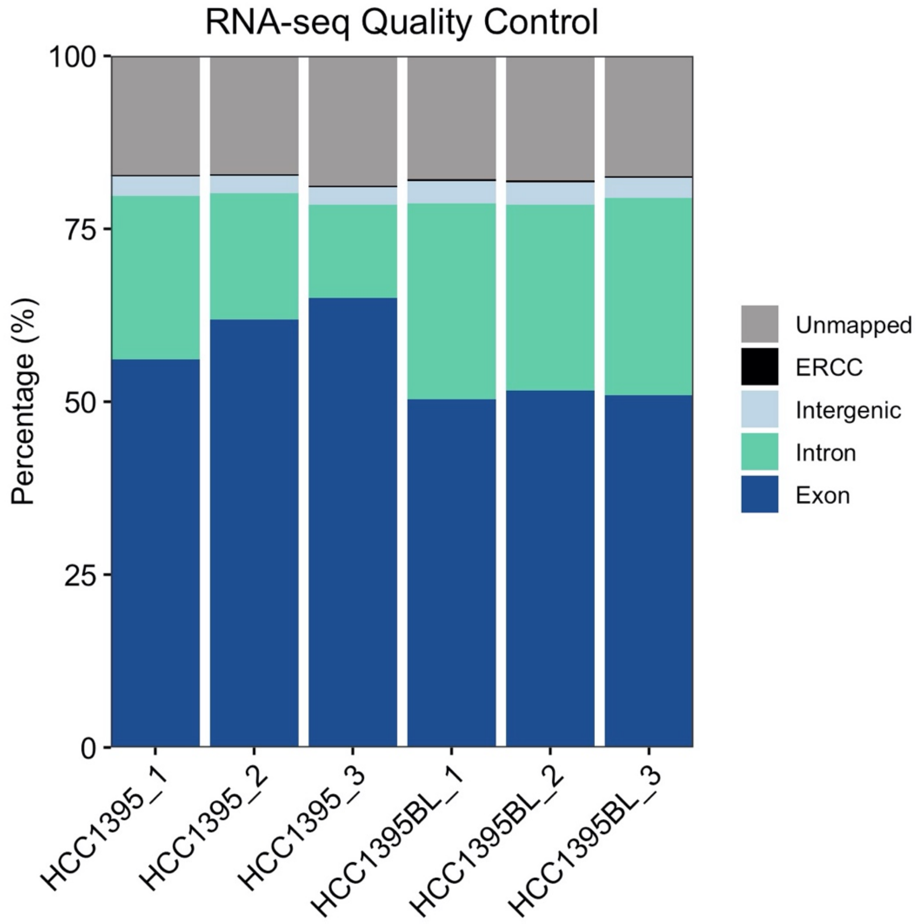


**Supplementary Figure 4. Mapping and alignment QC of bulk cell RNA-seq datasets.**

Bulk cell RNA-seq data sets were generated for both the HCC1395 and HCC1395BL cell lines (n=3 for each cell line). The figure shows the percentage of reads mapped to the exonic (dark blue), intronic (light green), intergenic (light blue) regions, Spiked in ERCC (1%) sequences (black), or reads not mapped to the human genome (gray) in the bulk RNA-seq data. The preprocessing pipeline for bulk RNA-seq data included QC (FastQC v0.11.4), trimming and filtering (Trimmomatic), alignment (STAR), and gene counting (RSEM).

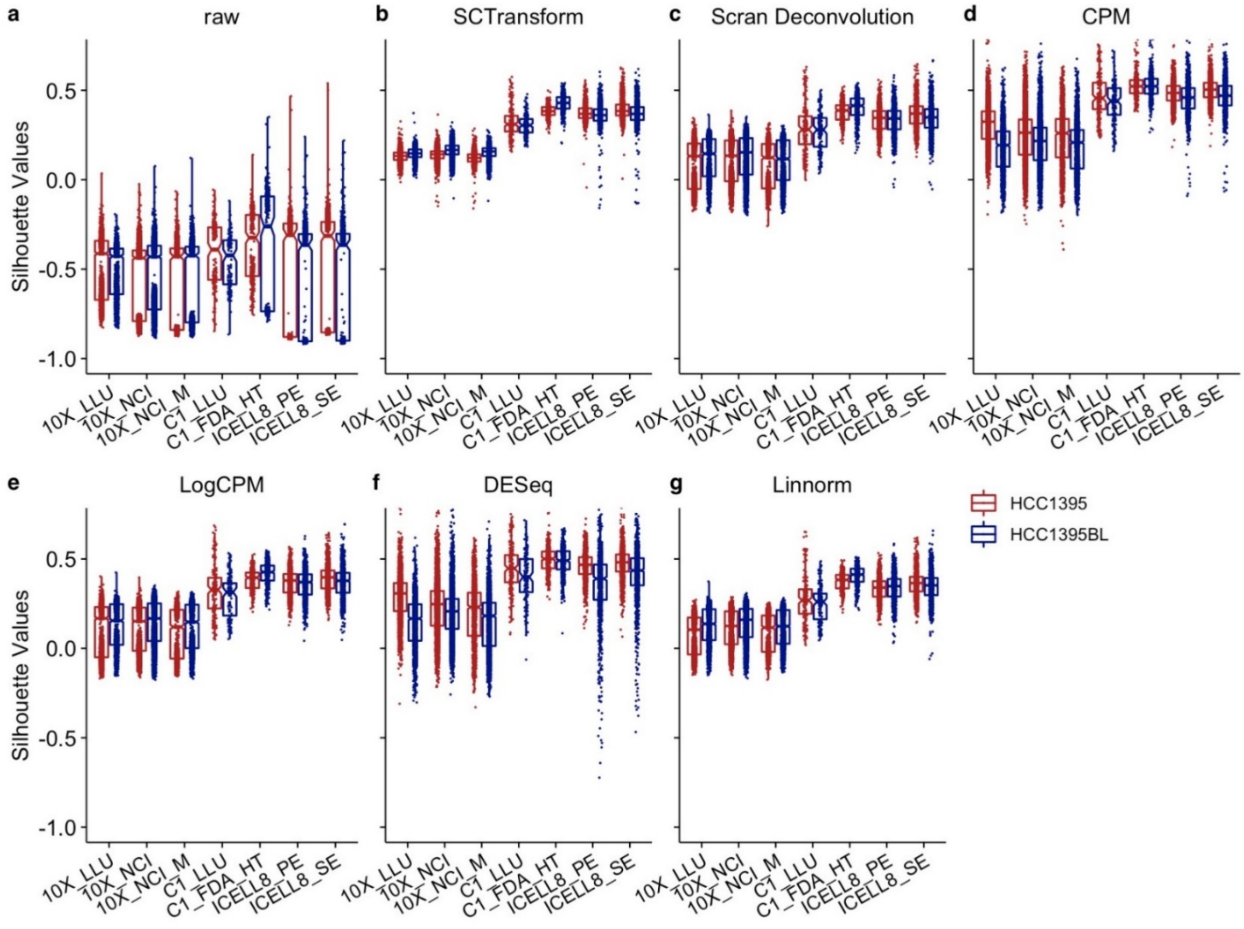


**Supplementary Figure 5. Comparison of silhouette scores across different scRNA-seq datasets.**

Fourteen scRNA-seq datasets were used to evaluate six normalization methods. **(a)** Boxplot of silhouette width values of raw gene expression across the different scRNA-seq platforms. **(b-g)** Boxplots of silhouette scores across the different scRNA-seq platforms using SCTransform **(b)**, scran deconvolution **(c)**, CPM **(d)**, LogCPM **(e)**, DESeq **(f)**, and Linnorm **(g)** normalizations. The scores for the C1 and ICELL8 platforms were consistently higher than the three 10X datasets.

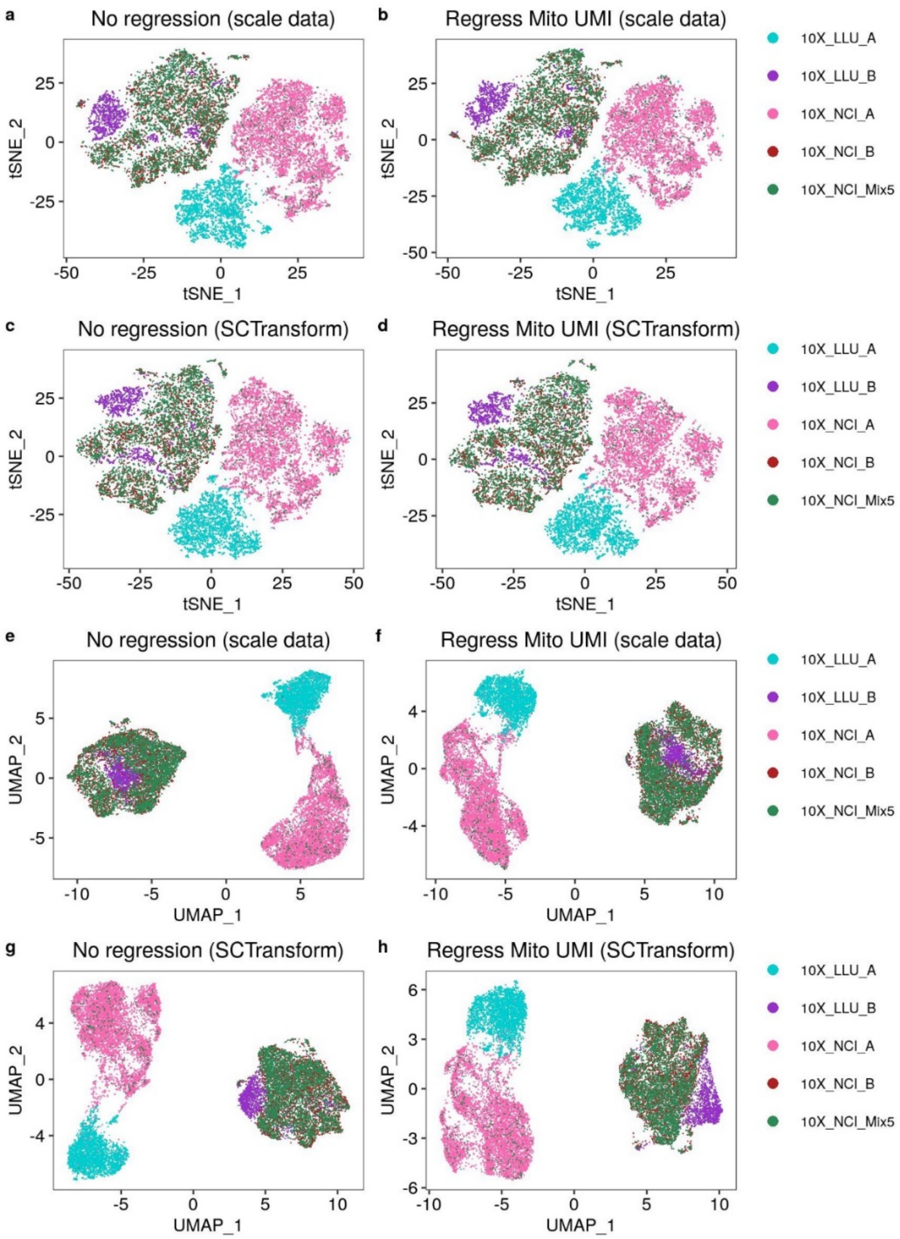


**Supplementary Figure 6. Regressing mitochondrial genes & normalizing UMI did not remove batch effects.**

**(a-d)** t-SNE and **(e-h)** UMAP plots generated using five samples prepared and sequenced at two sites after regressing out the effects of mitochondrial genes and UMI by Seurat v3. Five different batches of scRNA-seq data without mitochondrial gene regression and UMI normalization are shown as t-SNE **(a & c)** or UMAP plots **(e &g)**. After regression of mitochondrial genes (mito) and filtering cells with mito >5% and normalizing with either logNormalize or SCTransform, the data are shown as t-SNE **(b & d)** or UMAP plots **(f & h)**. Regressing the mitochondrial genes and normalizing the dataset did not remove the observed batch effects.

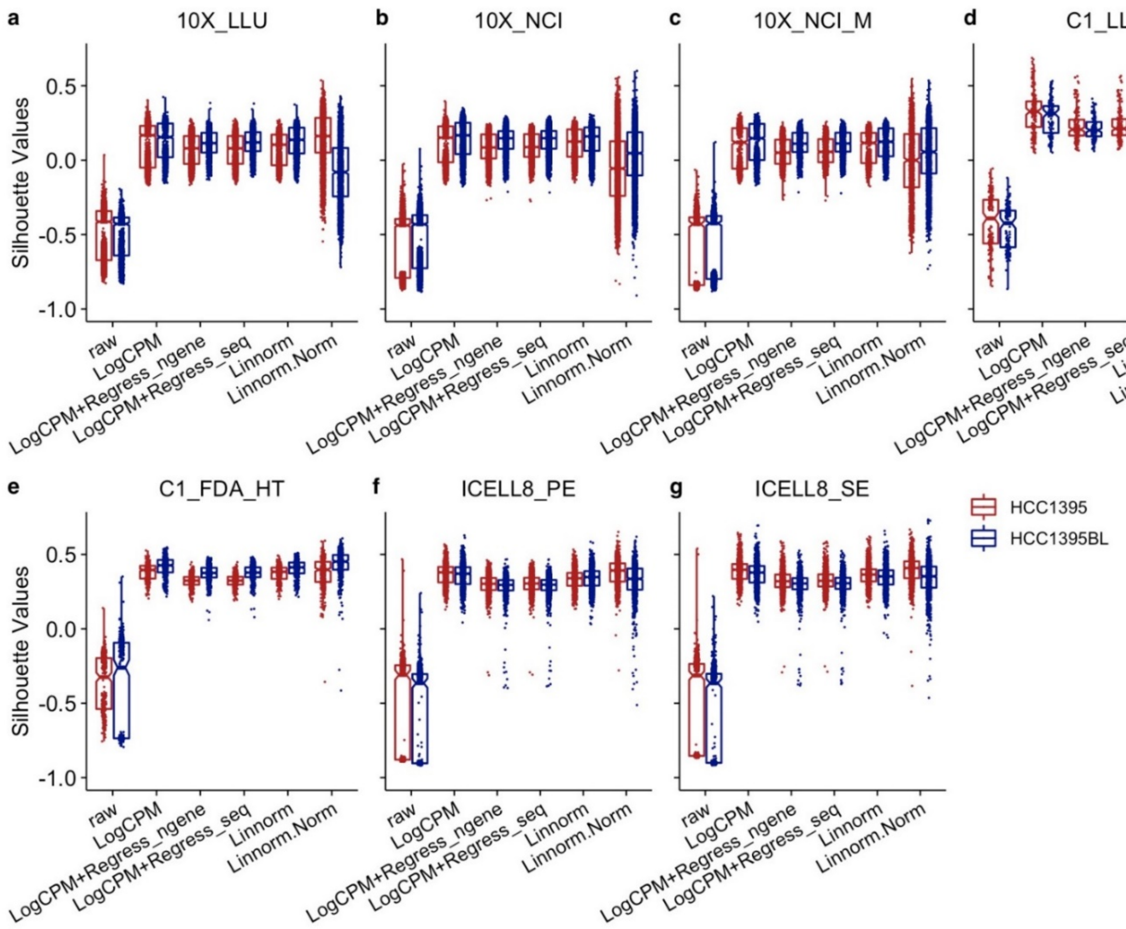


**Supplementary Figure 7. Regressing out number of genes detected did not improve the silhouette scores.**

**(a-g)** Boxplots of silhouette width values stratified by regression-based normalization methods, LogCPM and two different Linnorm methods across seven datasets and platforms: (**a)** 10X_LLU, (**b**) 10X_NCI, (**c**) 10X_NCI_M, (**d**) C1_LLU, (**e**) C1_FDA_HT, (**f**) ICELL8_PE, (**g**) ICELL8_SE. Each dot represents a cell. X-axis represents the normalization methods; Y-axis represents the silhouette width values for every pair of cells.

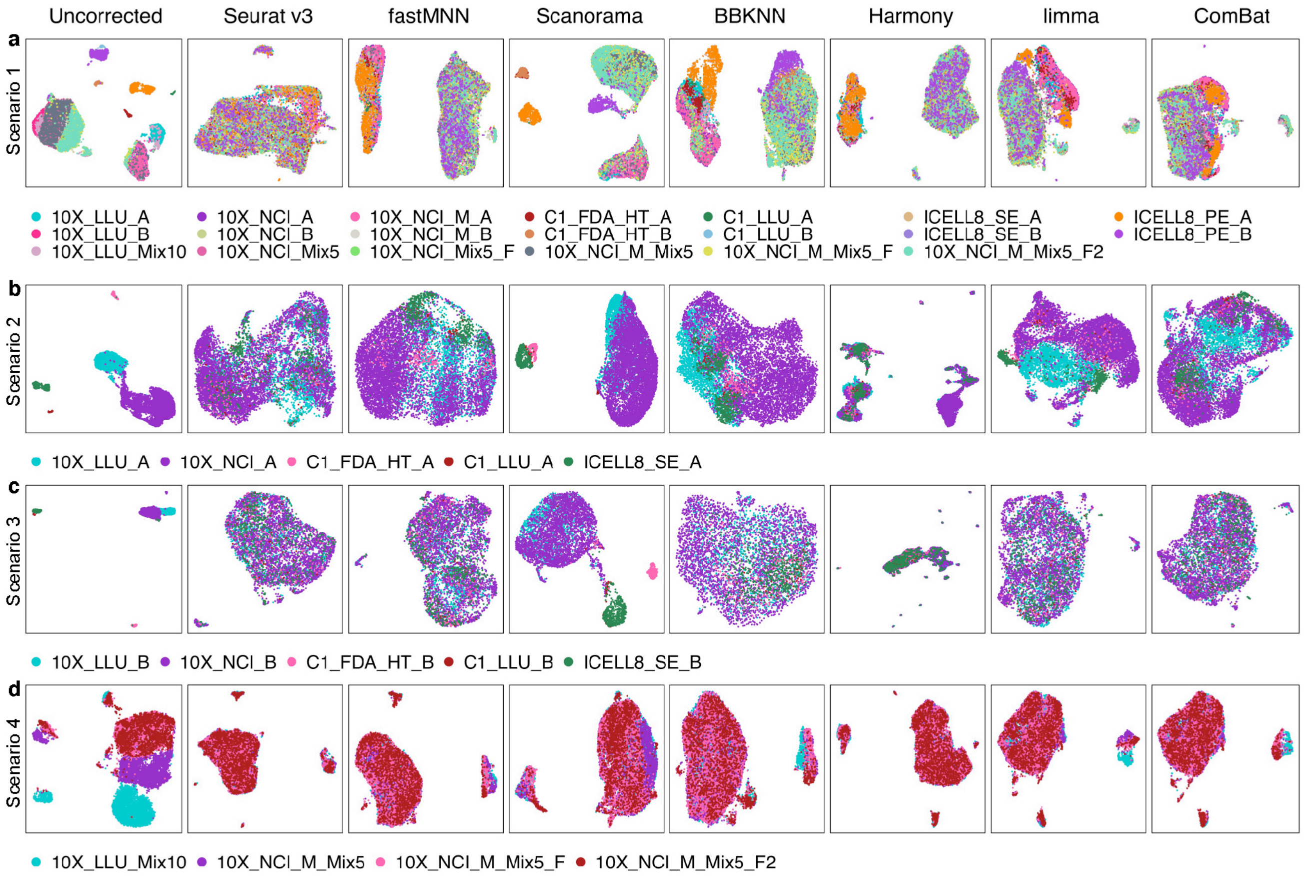


**Supplementary Figure 8. UMAP projection showing batch effect correction by mixability and clusterability using scRNA-seq datasets in four different sample scenarios.**

**(a)** Batch effect corrections performed using 20 scRNA-seq datasets across different sites and platforms. Batch correction methods included Seurat v3.1, fastMNN (SeuratWrappers v0.1.0), Scanorama V1.4, BBKNN V1.3.5, Harmony V0.99.9, limma V3.40.4, and Combat (sva V3.32.1). **(b-c)** Batch-effect corrections performed using five scRNA-seq datasets from different sites and/or platforms in biologically similar cells, either breast cancer cells **(b)** or B lymphocytes **(c)**. The five scRNA-seq datasets from different platforms were: 10X_LLU, C1_FDA_HT, 10X_NCI, C1_LLU, and ICELL8_SE. **(d)** Batch-effect corrections performed using scRNA-seq datasets derived from spiked-in mixtures of cells in which either 5% or 10% cancer cells were spiked into the sample B cells. Four datasets were analyzed: 10X_LLU_Mix10, 10X_NCI_M_Mix5, 10X_NCI_M_Mix5_F, 10X_NCI_M_Mix5_F2. The top 2000 highly variable genes (HVGs) of these datasets were used as the gene set for batch correction. All the 10X data were preprocessed using CellRanger 3.1.

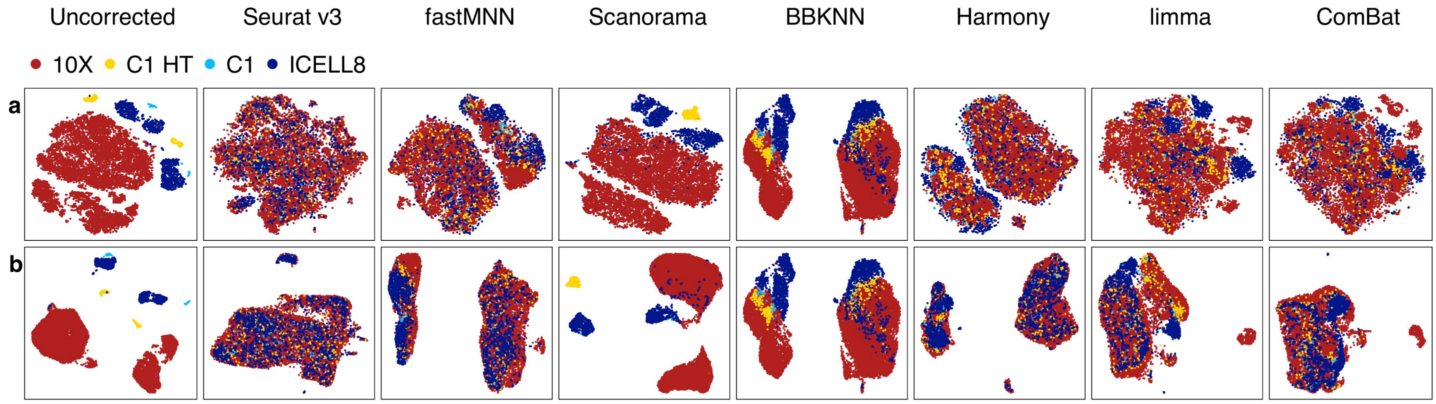


**Supplementary Figure 9. t-SNE and UMAP projections showing batch effect corrections by mixability and clusterability across four scRNA-seq platforms.**

**(a)** tSNE and **(b)** UMAP plots showing the batch-effect corrections performed by seven methods using 20 scRNA-seq datasets across different platforms and sites. Note, for BBKNN, only UMAP projections were available and are shown in both (**a**) and (**b**). The pair-wise breast cancer cells and B cells captured from different scRNA-seq platforms were labeled in different colors: 10X 3' scRNA-seq platform (red color), C1 3' HT scRNA-seq platform (yellow color), C1 full-length scRNA-seq platform (light blue color), ICELL8 full-length scRNA-seq platform (dark blue color). Batch correction methods included: Seurat v3.1, fastMNN (SeuratWrappers v0.1.0), Scanorama V1.4, BBKNN V1.3.5, Harmony V0.99.9, limma V3.40.4, and Combat (sva V3.32.1). Scanorama was unable to correct batch effects when non-10X platforms were included in the analysis. The top 2000 HVGs across all data sets were used as the gene set for batch correction. All the 10X data were preprocessed using CellRanger 3.1.

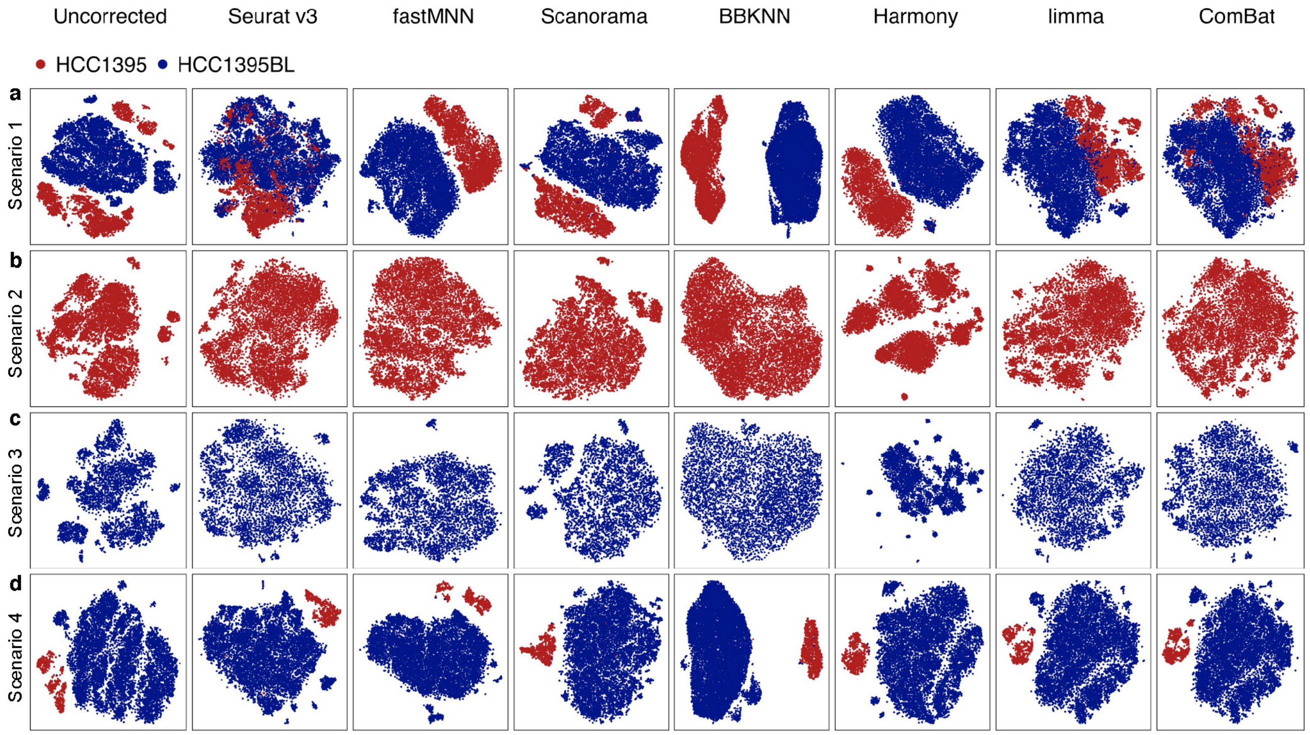


**Supplementary Figure 10. Batch effect correction displayed by cell type identity.**

**(a)** Batch-effect corrections performed using 20 scRNA-seq datasets across different sites and platforms. **(b)-(c)** Batch-effect corrections performed using five scRNA-seq datasets from different sites and/or platforms in biologically similar cells, either breast cancer cells **(b)** or B lymphocytes **(c)**. The five scRNA-seq dataset were: 10X_LLU, C1_FDA_HT, 10X_NCI, C1_LLU, and ICELL8_SE. **(d)** Batch-effect corrections performed using scRNA-seq data derived from spiked-in mixtures of cells in which either 5% or 10% cancer cells were spiked into the sample B cells. Four datasets were analyzed: 10X_LLU_Mix10, 10X_NCI_M_Mix5, 10X_NCI_M_Mix5_F, 10X_NCI_M_Mix5_F2. For BBKNN, only UMAP projections were available and shown in (**a-d**). The HCC1395 breast cancer cells (A) were labeled in red color and the HCC1395BL B lymphocytes (B) were labeled in blue color. Batch correction methods included Seurat v3.1, fastMNN (SeuratWrappers v0.1.0), Scanorama V1.4, BBKNN V1.3.5, Harmony V0.99.9, limma V3.40.4, and Combat (sva V3.32.1). The top 2000 HVGs were used as the gene set for batch correction. All the 10X data were preprocessed using CellRanger 3.1.

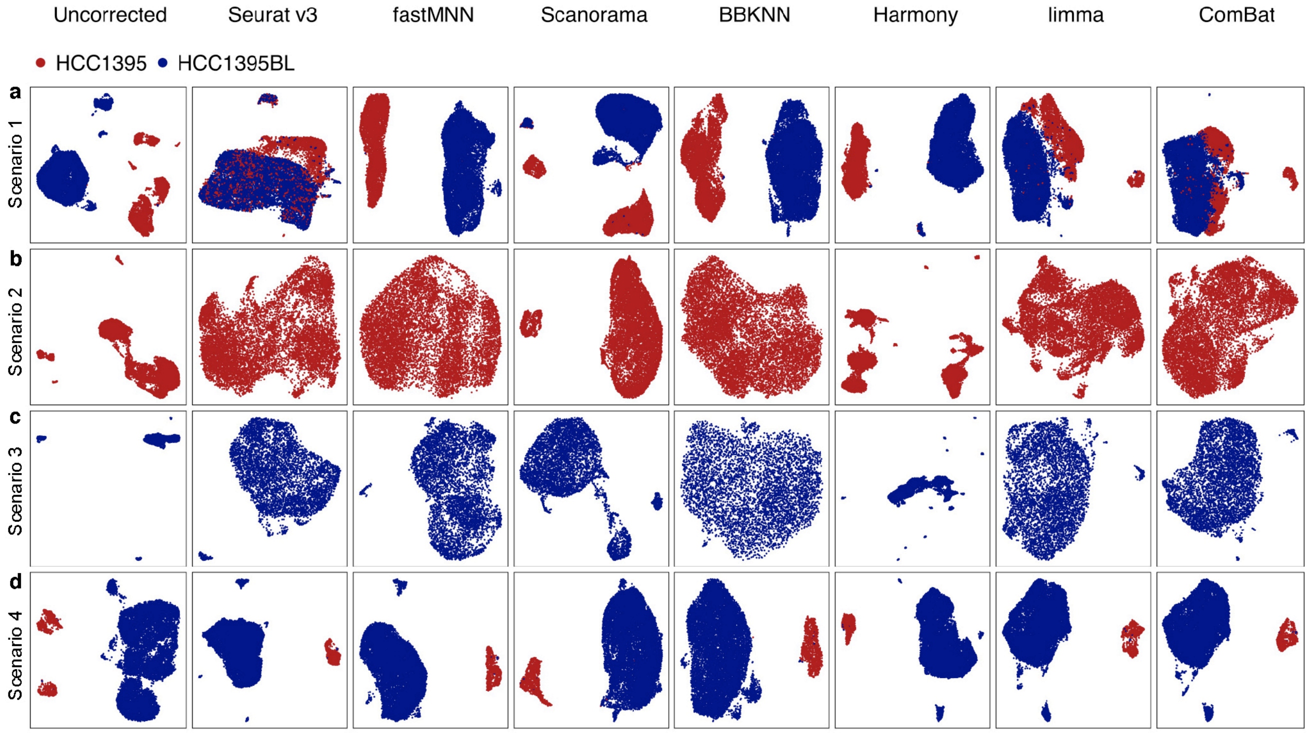


**Supplementary Figure 11. UMAP projection showing batch-effect correction displayed according to cell-type identity.**

**(a)** Batch-effect corrections performed using 20 scRNA-seq datasets across different sites and platforms. **(b-c)** Batch-effect corrections performed using five scRNA-seq datasets from different sites and/or platforms in biologically similar cells, either breast cancer cells **(b)** or B lymphocytes **(c)**. The five scRNA-seq dataset were: 10X_LLU, C1_FDA_HT, 10X_NCI, C1_LLU, and ICELL8_SE. **(d)** Batch-effect corrections performed using scRNA-seq data derived from spiked-in mixtures of cells in which either 5% or 10% cancer cells were spiked into the sample B cells. Four datasets were analyzed: 10X_LLU_Mix10, 10X_NCI_M_Mix5, 10X_NCI_M_Mix5_F, 10X_NCI_M_Mix5_F2. The HCC1395 breast cancer cells (A) were labeled in red color and the HCC1395BL B lymphocytes (B) were labeled in blue color. Batch correction methods included Seurat v3.1, fastMNN (SeuratWrappers v0.1.0), Scanorama V1.4, BBKNN V1.3.5, Harmony V0.99.9, limma V3.40.4, and Combat (sva V3.32.1). The top 2000 HVGs were used as the gene set for batch correction. All the 10X data were preprocessed using CellRanger 3.1.

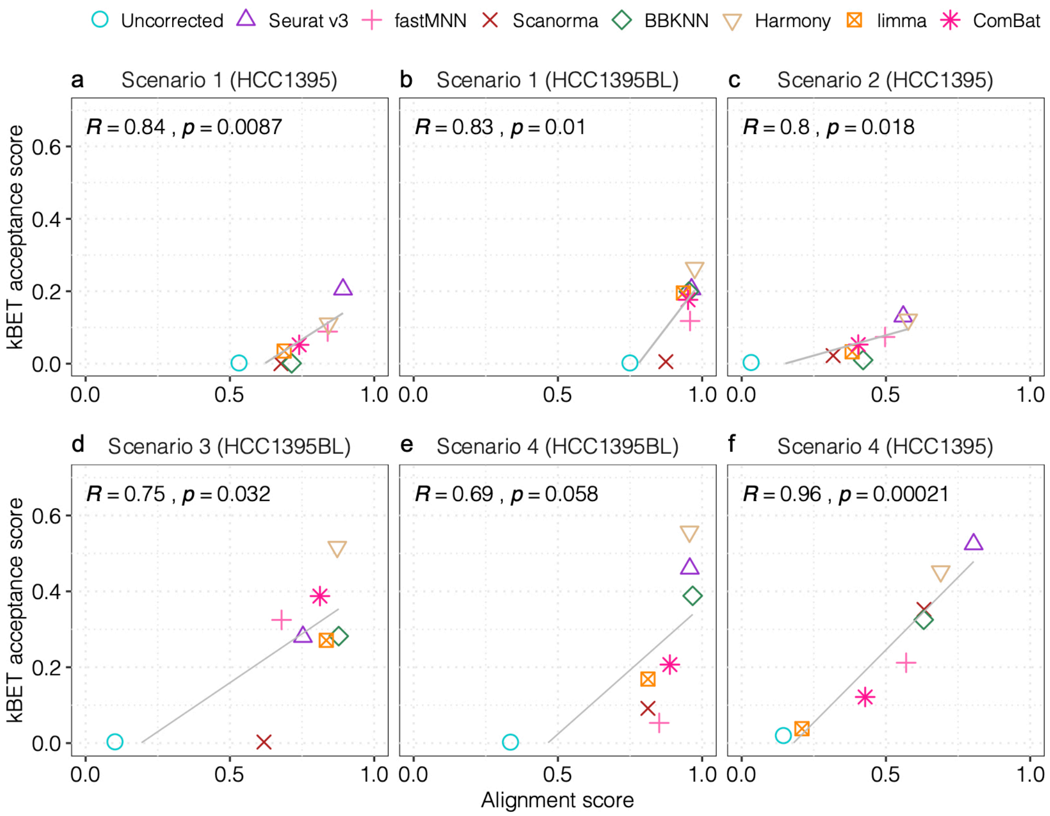


**Supplementary Figure 12. Correlation between kBET acceptance score and modified alignment score.**

**(a-b)** Modified alignment and kBET acceptance scores were calculated using 20 scRNA-seq datasets (scenario #1). The scores were calculated for breast cancer cells **(a)** and B lymphocytes **(b)**, respectively. **(c-d)** Modified alignment and kBET scores were calculated using biologically similar samples (scenario #2), either breast cancer cells **(c)** or B lymphocytes **(d)**. **(e-f)** Modified alignment and kBET scores were calculated using scRNA-seq data derived from spiked-in mixture samples in which either 5% or 10% cancer cells were spiked into the B lymphocytes. The scores were calculated for B lymphocytes **(e)** or breast cancer cells **(f)**, respectively. Correlation (R) between the alignment score and kBET acceptance score was shown in each plot along with P value.

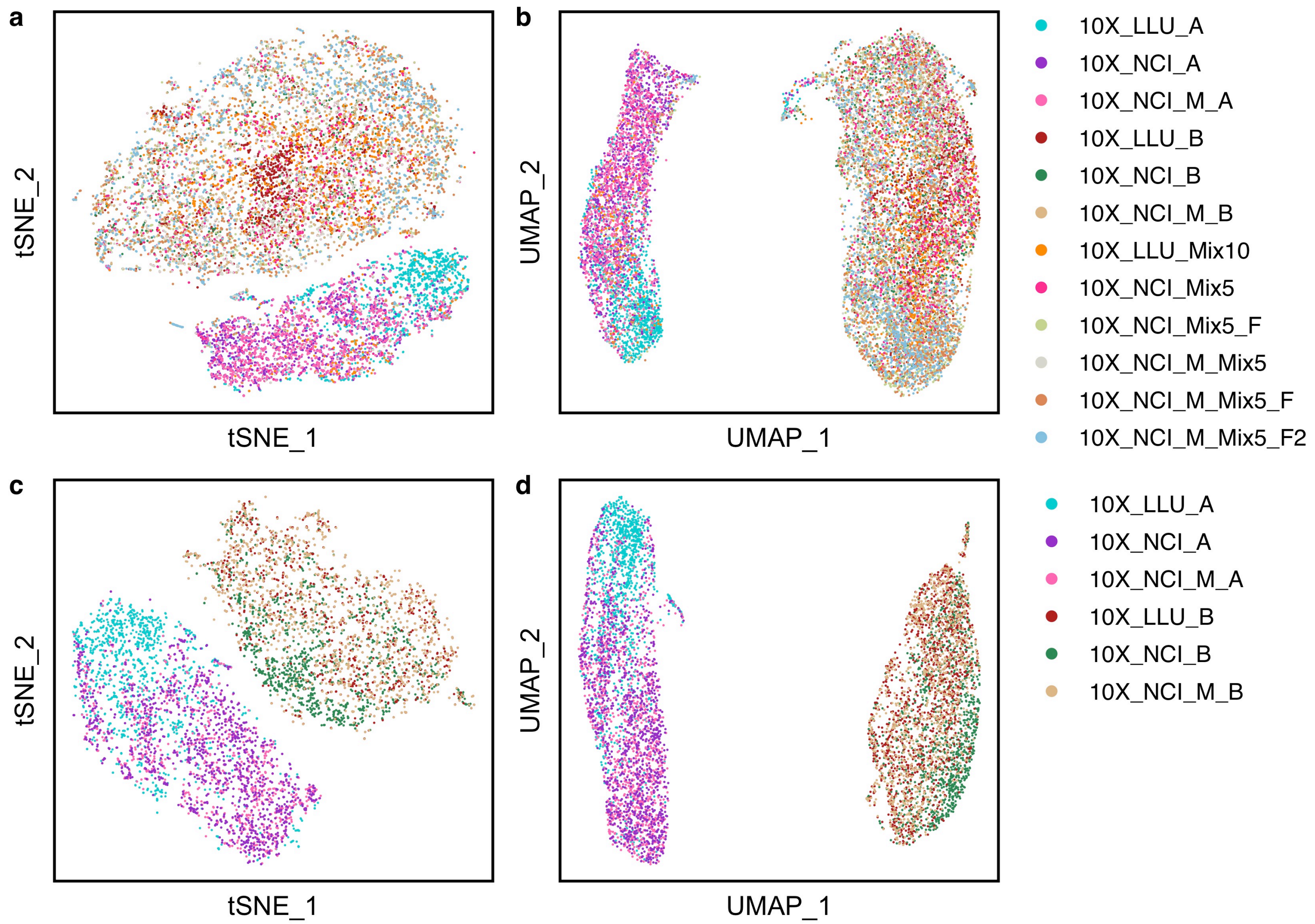


**Supplementary Figure 13. Scanorama worked well for 10X Genomics scRNA-seq datasets regardless of the presence of shared cell types across batches.**

**(a)** t-SNE and **(b)** UMAP plots showing projections of batch-effect corrections using twelve 10X Genomics scRNA-seq datasets consisting of both mixed and non-mixed samples from two sites in different batches after Scanorama (version 1.4.) batch correction. **(c)** t-SNE and **(d)** UMAP plots showing projections of batch-effect corrections using six 10X Genomics scRNA-seq datasets consisting of only non-mixed samples from two sites in different batches after Scanorama (version 1.4.) batch correction. Different colors represent different datasets. All the datasets were subsampled to 1200 cells. After the batch correction, cells from the same cell line type tended to cluster together and mixed well within the same cell types. All the data were preprocessed using CellRanger 3.1.

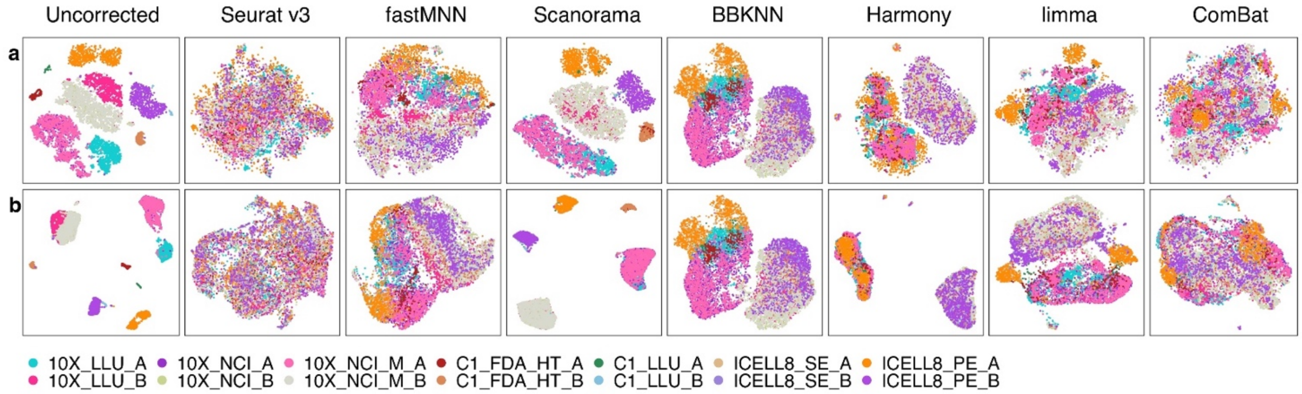


**Supplementary Figure 14. Batch-effect correction evaluating clusterability using 14 scRNA-seq datasets without spiked-in mixtures.**

**(a)** t-SNE and **(b)** UMAP plots showing batch-effect corrections performed by seven methods using 14 scRNA-seq non-mixture datasets across different platforms and sites. Note, for BBKNN, only UMAP projections were available and shown in (**a-b)**. Six spiked-in mixture scRNA-seq datasets (10X_LLU_Mix10, 10X_NCI_Mix5, 10X_NCI_Mix5_F, 10X_NCI_M_Mix5, 10X_NCI_M_Mix5_F, and 10X_NCI_M_Mix5_F2) were removed from the 20 datasets in scenario 1 for batch-effect correction evaluation. Batch correction methods included Seurat v3.1, fastMNN (SeuratWrappers v0.1.0), Scanorama V1.4, BBKNN V1.3.5, Harmony V0.99.9, limma V3.40.4, and Combat (sva V3.32.1). All the 10X data were preprocessed using CellRanger 3.1.

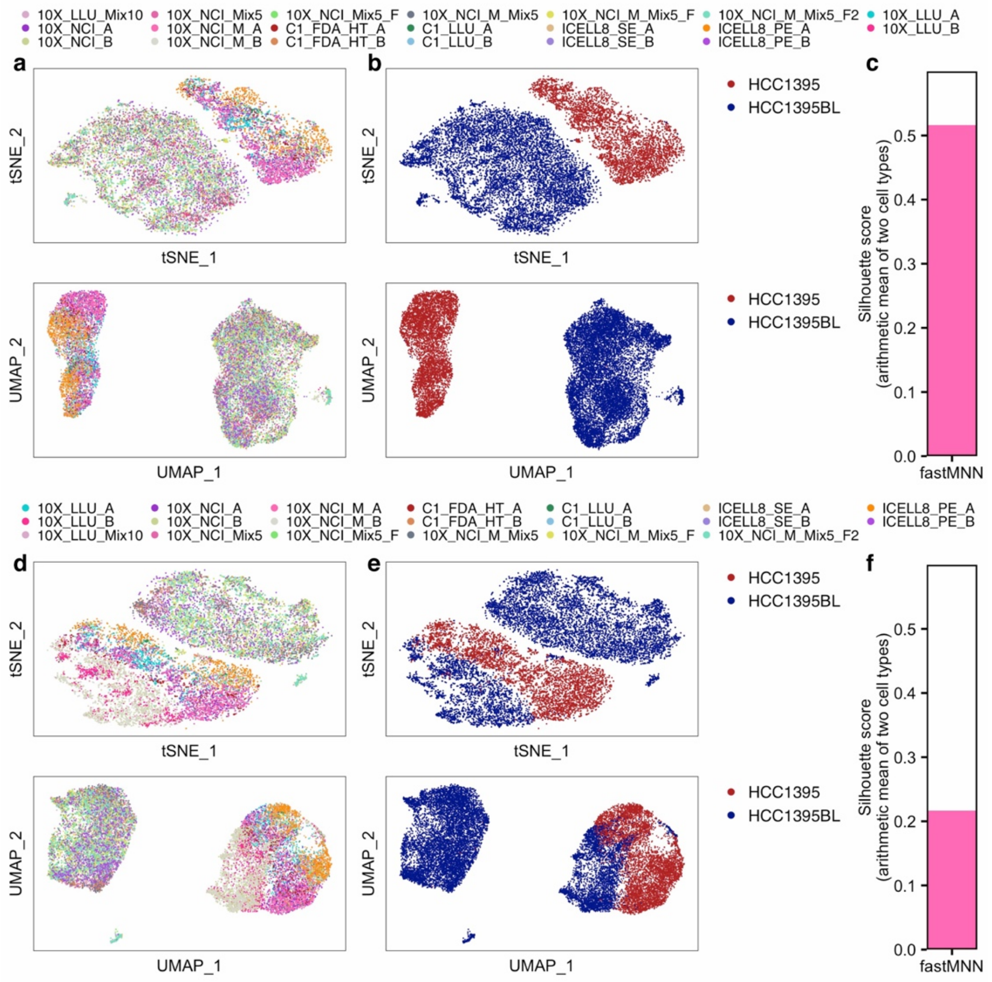


**Supplementary Figure 15. fastMNN batch-effect correction depends on the order of importing scRNA-seq data into the pipeline**.

**(a)** t-SNE and **(b)** UMAP projections showing that fastMNN correctly separated the two cell types into two clusters of either cancer cells (red) or B cells (blue). **(c)** A *silhouette* score = 0.52 was obtained when the spiked-in datasets were imported into the pipeline before the non-mixed scRNA-seq datasets. Both batch-effect correction and silhouette width calculation were performed using fastMNN (SeuratWrappers v0.1.0), when the spiked-in (mixed) datasets (i.e., 10X_LLU_Mix10, 10X_NCI_Mix5, 10X_NCI_Mix5_F, 10X_NCI_M_Mix5, 10X_NCI_M_Mix5_F and 10X_NCI_M_Mix5_F2) were imported into the pipeline before other non-mixed scRNA-seq datasets using the *silhouette* function from the R package *cluster* (v.2.0.8) and fastMNN embeddings file. **(d-e)** Batch-effect correction performed using fastMNN (SeuratWrappers v0.1.0) when the non-mixed datasets were imported into the pipeline before the mixture datasets. **(f)** A low *silhouette* score of 0.22 obtained when the spiked-in datasets were imported into the pipeline after the non-mixed scRNA-seq datasets. Twenty scRNA-seq datasets were included for evaluation. All the datasets were subsampled to 1200 cells. The order of dataset input is shown on the top of the figure **(a-c** or **d-f)**.

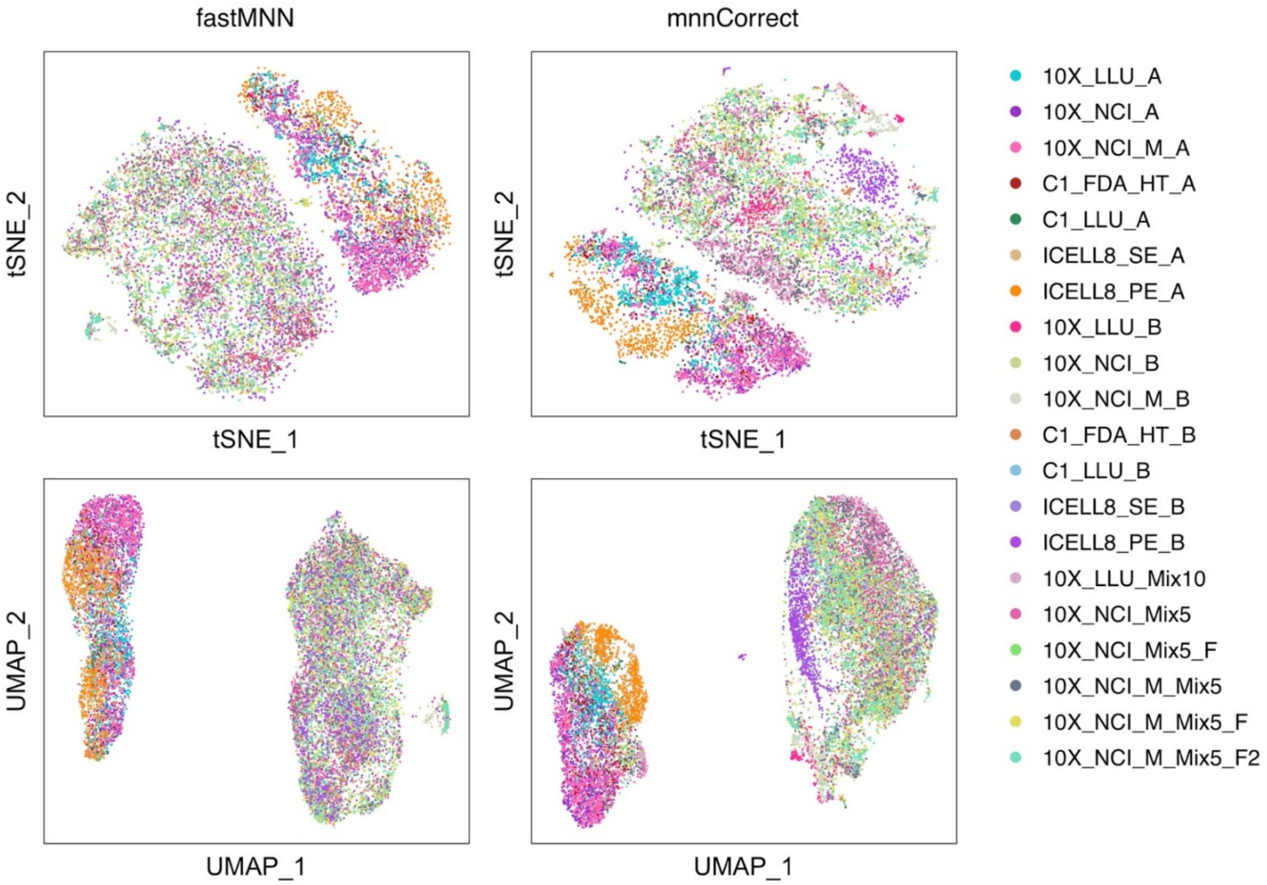


**Supplementary Figure 16. Batch-effect correction comparison between MNN and fastMNN using 20 scRNA-seq datasets.**

Twenty scRNA-seq datasets were used for evaluation. All the datasets were subsampled to 1200 cells. The same package versions were used as indicated in the Fig. 4 and Supplementary Figure 8. The samples were loaded into the pipeline in the same order as in the Suppl. Figure 15a-b. The two methods worked similarly except that fastMNN used significantly less time (51.1 seconds) compared with mnnCorrect (62781.8 seconds). Both methods were compared using the same computer system: DELL PRECISION TOWER 7910, 128G RAM (Memory), 14 CPU cores (56 CPU processors), Intel, Xeon CPU, E5-2680 V4@2.40Ghz.

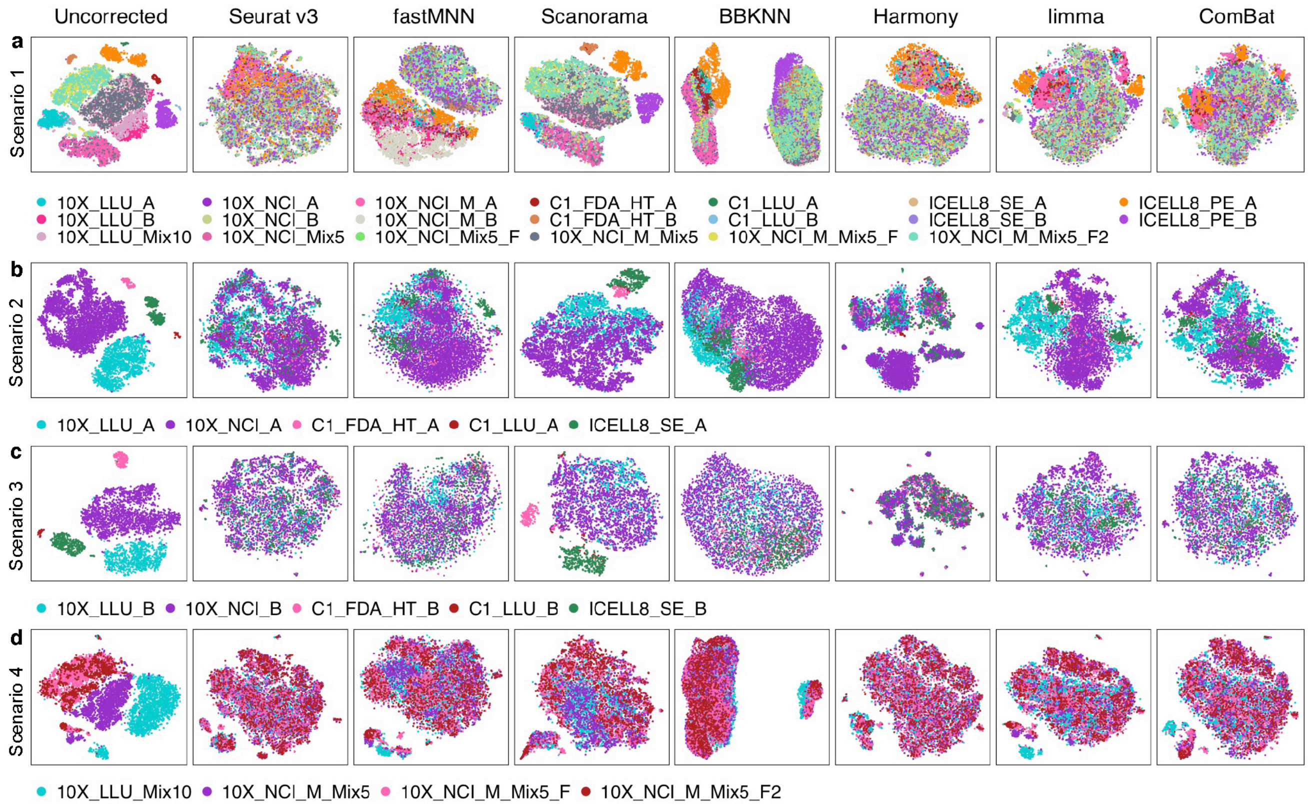


**Supplementary Figure 17. t-SNE projection showing batch-effect correction using CellRanger 2.0 preprocessed 10X data.**

**(a)** Batch-effect corrections performed using 20 scRNA-seq datasets across different sites and platforms. **(b-c)** Batch-effect corrections performed using five scRNA-seq datasets from different sites and/or platforms in biologically similar cells, either breast cancer cells **(b)** or B lymphocytes **(c)**. The five scRNA-seq datasets were from: 10X_LLU, C1_FDA_HT, 10X_NCI, C1_LLU, and ICELL8_SE. **(d)** Batch-effect corrections performed using 10X scRNA-seq data from two centers derived from spiked-in mixtures of cells in which either 5% or 10% cancer cells were spiked into the sample B cells. Four datasets were analyzed (**d**): 10X_LLU_Mix10, 10X_NCI_M_Mix5, 10X_NCI_M_Mix5_F, 10X_NCI_M_Mix5_F2. Batch correction methods included: Seurat v3.1, fastMNN (SeuratWrappers v0.1.0), Scanorama V1.4, BBKNN V1.3.5, Harmony V0.99.9, limma V3.40.4, and Combat (sva V3.32.1). For BBKNN, only UMAP projections were available and shown in (**a-d)**. The top 2000 HVGs were used as the gene set for batch correction. All the 10X data were preprocessed using CellRanger 2.0. Note, for BBKNN, only UMAP projections were available.

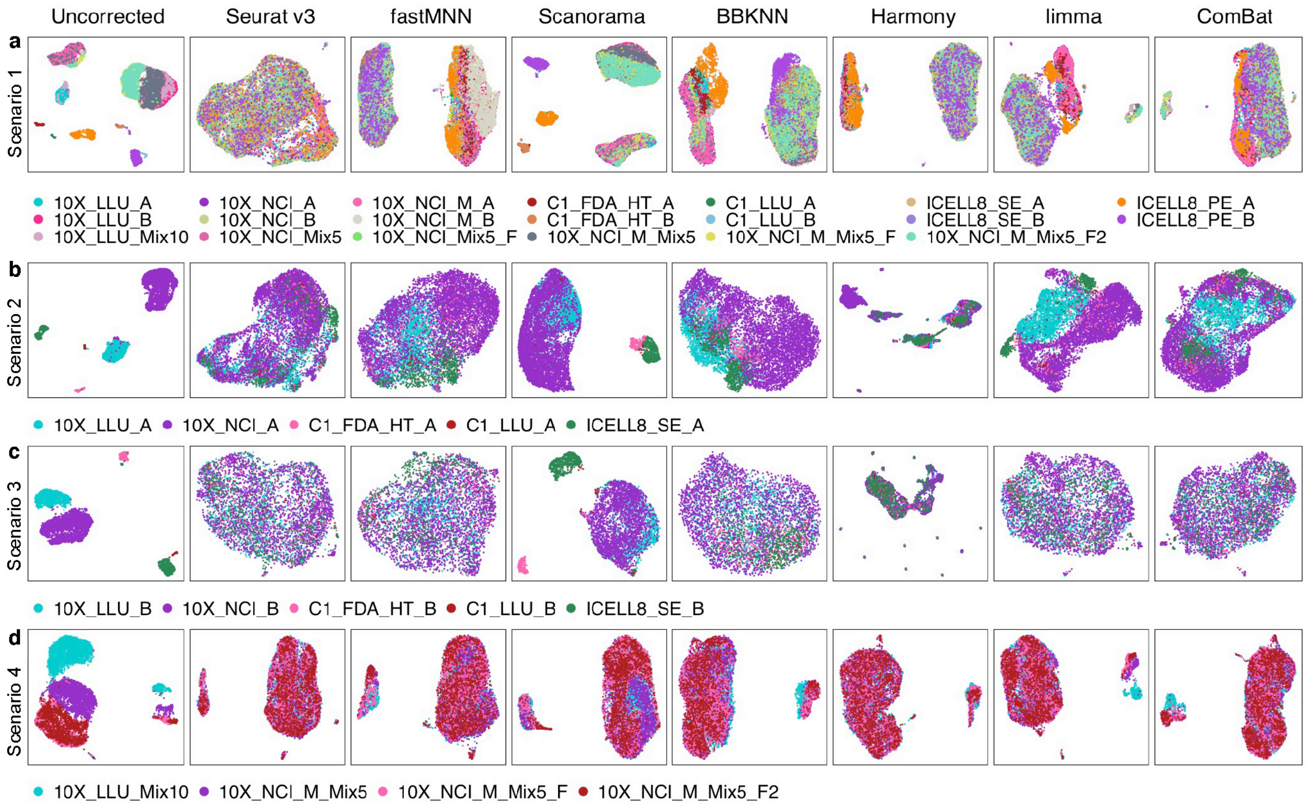


**Supplementary Figure 18. UMAP projection showing batch-effect correction using CellRanger 2.0 preprocessed 10X data.**

**(a)** Batch-effect corrections performed using 20 scRNA-seq datasets across different sites and platforms. **(b-c)** Batch-effect corrections performed using 5 scRNA-seq datasets from different sites and/or platforms in biologically similar cells, either breast cancer cells **(b)** or B lymphocytes **(c)**. The five scRNA-seq datasets were from: 10X_LLU, C1_FDA_HT, 10X_NCI, C1_LLU, and ICELL8_SE. **(d)** Batch-effect corrections performed using 10X scRNA-seq data from two centers derived from spiked-in mixtures of cells in which either 5% or 10% cancer cells were spiked into the sample B cells. Four datasets were analyzed (**d**): 10X_LLU_Mix10, 10X_NCI_M_Mix5, 10X_NCI_M_Mix5_F, 10X_NCI_M_Mix5_F2. Batch correction methods included: Seurat v3.1, fastMNN (SeuratWrappers v0.1.0), Scanorama V1.4, BBKNN V1.3.5, Harmony V0.99.9, limma V3.40.4, and Combat (sva V3.32.1). The top 2000 HVGs were used as the gene set for batch correction. All the 10X data were preprocessed using CellRanger 2.0.

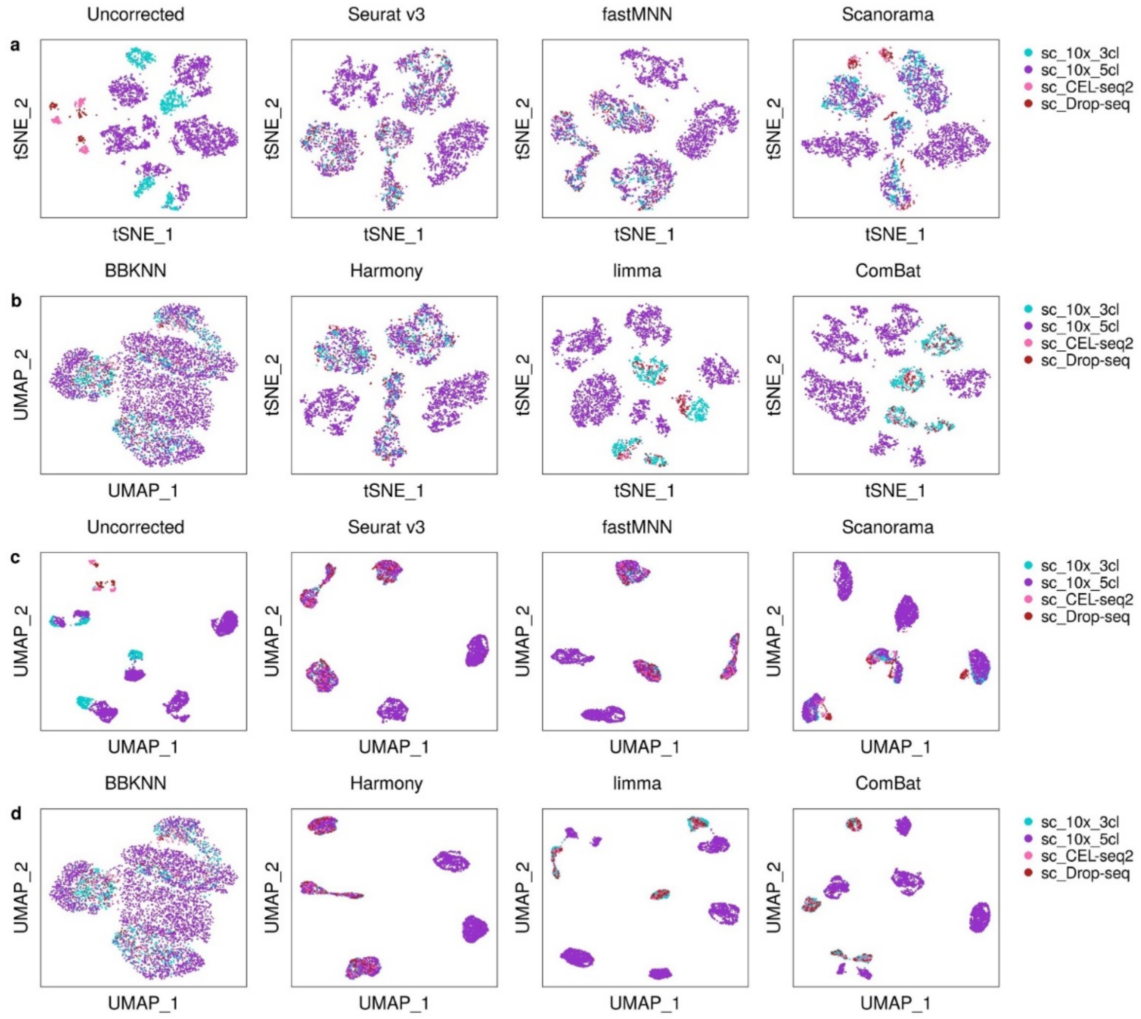


**Supplementary Figure 19. Batch-effect correction using Tian's scRNA-seq data preprocessed using CellRanger v2.0 and umitools v1.0.0.**

**(a-b)** t-SNE and **(c-d)** UMAP plots generated using four scRNA-seq datasets from Tian et al. and 7 batch correction methods. For BBKNN, only UMAP projections were available and shown in (**b** & **d**). Four datasets derived from two experimental batches were: 10X scRNA-seq data from three mixed lung cancer cell lines, 10X scRNA-seq data from five mixed lung cancer cell lines, CELL-seq2 data from three mixed lung cancer cell lines, and Drop-seq data from three mixed lung cancer cell lines. The two 10X datasets were preprocessed with CellRanger 2.0. The other two datasets were pre-processed by umitools v1.0.0. The subsequent batch-effect corrections were carried out using the same methods that were applied to our data as described in the Methods section.

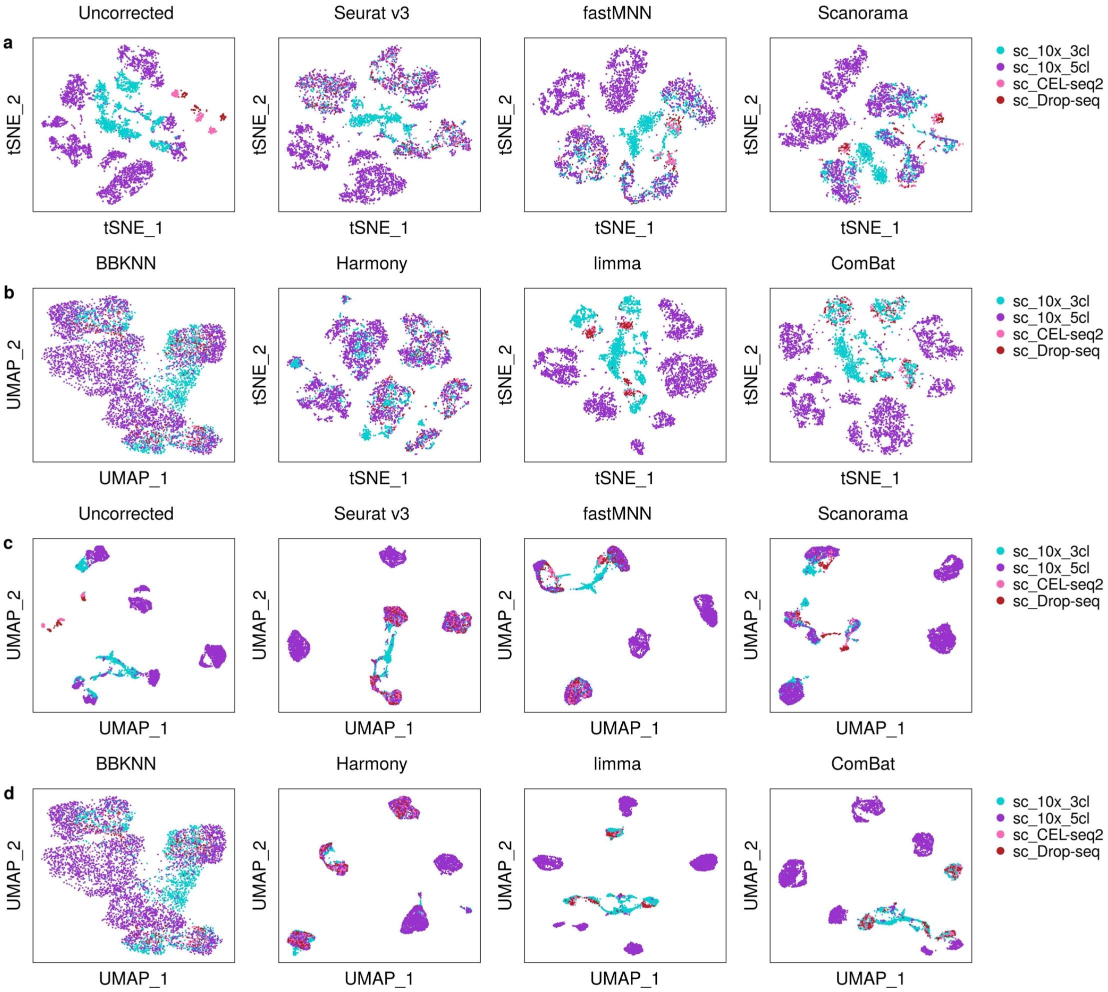


**Supplementary Figure 20. Batch-effect correction using Tian's scRNA-seq data preprocessed using CellRanger v3.0.2 and umitools v1.0.0.**

**(a-b)** t-SNE and **(c-d)** UMAP plots generated using four scRNA-seq datasets from Tian et al. and 7 batch correction methods. For BBKNN, only UMAP projections were available and shown in (**b & d**). The four scRNA-seq datasets were derived from two experimental batches, i.e., 10X data, CELL-seq2 data, and Drop-seq data (each made using a mixture of three lung cancer cell lines) in the 1$^{st}$ batch; and 10X data from a mixture of five lung cancer cell lines in the 2$^{nd}$ batch. The 10x datasets were preprocessed using CellRanger 3.0.2. The other two datasets were pre-processed by umitools v1.0.0. The subsequent batch-effect corrections were carried out using the same methods that were applied to our data, as described in the Methods section.

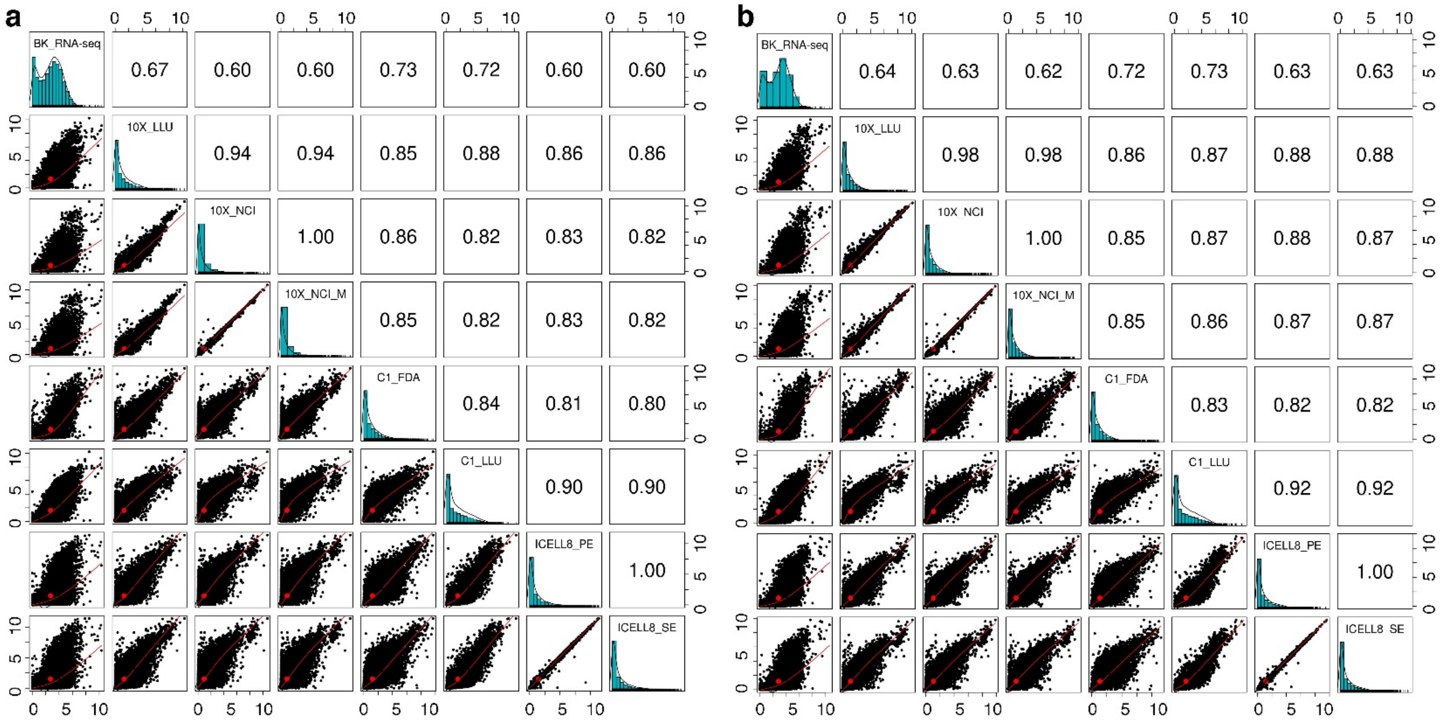


**Supplementary Figure 21. Scatter plots of gene expression profiles across seven scRNA-seq datasets.**

The scatter plots displaying the correlation of gene expression profiles between each pair of datasets for either breast cancer cells (**a**) or B lymphocytes (**b**). The commonly detected transcripts [(log(CPM +1) normalized] across all datasets were used (15,553 genes for breast cancer cells and 15,201 genes for B lymphocytes) to generate the scatter plots. Each dot represents each gene as a point in each scatterplot; $x, y$ values represent the gene expression variation in a pair of compared datasets. The middle diagonal bar charts display the distribution of the most abundant or rare genes in each scRNA-seq dataset or platform. The Pearson correlation between each of the compared datasets is shown on the top panel of the plot to show the consistency of the different RNA-seq datasets.

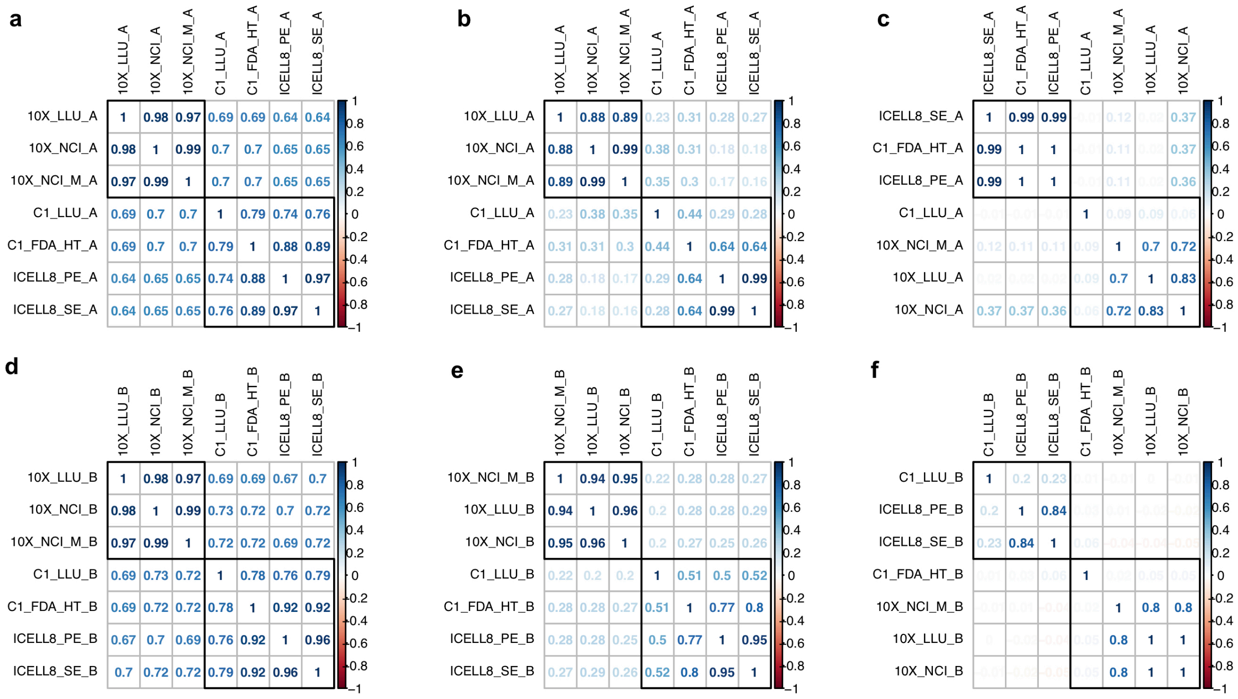

**Supplementary Figure 22. Pearson correlation coefficients across platforms and centers based on gene expression levels.**

**(a-f)** Benchmarking consistency of highly abundant, intermediate, and low-abundance genes across all seven scRNA-seq datasets. Data shown for cancer cells (**a**-**c**) and B cells (**d**-**f**). Pairwise Pearson correlation of the percentage of cells which expressed the 500 highly-expressed genes (**a & d)**; 500 intermediately expressed genes **(b & e)**; and 500 low-abundance genes **(c & f)**. To get the comparable cell percentages, we only considered gene count matrices from the down-sampling results (100K reads per cell) of zUMIs (10X datasets) and featureCounts (non-10X datasets) pipelines. The Pearson correlations of the cell percentage between any two scRNA-seq datasets were calculated for each of the three expression groups to evaluate the consistency.

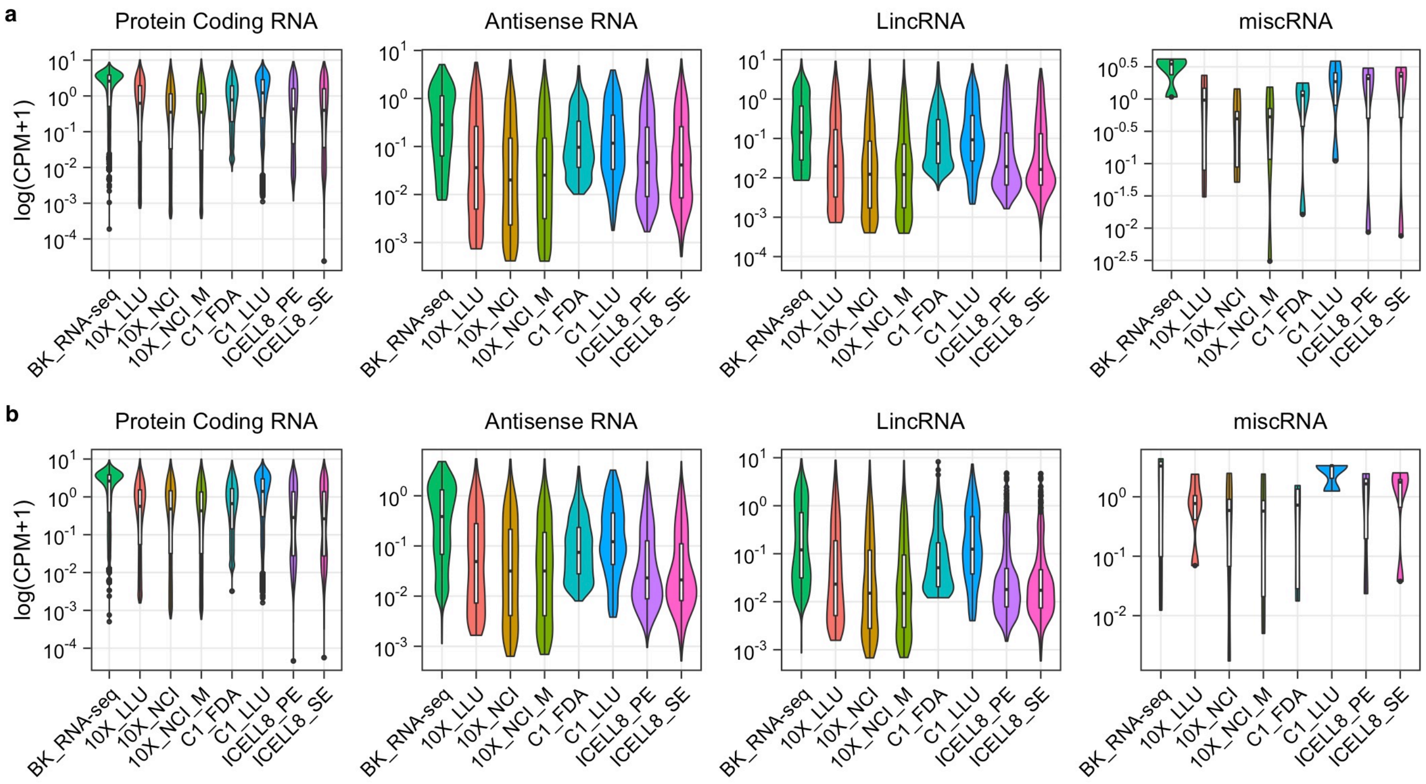


**Supplementary Figure 23. Consistency of protein coding, antisense, LincRNA, and miscRNA gene expression across seven scRNA-seq datasets.**

Violin plots generated based on gene expression profiles for **(a)** breast cancer cells, and **(b)** B lymphocytes in the selected four RNA categories across eight different datasets. The raw gene count matrices were converted to a normalized gene list using the method described in the Methods section. The datasets included bulk cell RNA-seq (BK_RNA-seq), 3' scRNA-seq (10X_LLU, 10X_NCI, 10X_NCI_M, C1_FDA_HT), and full-length transcripts (C1_LLU, ICELL8_PE and ICELL8_SE).

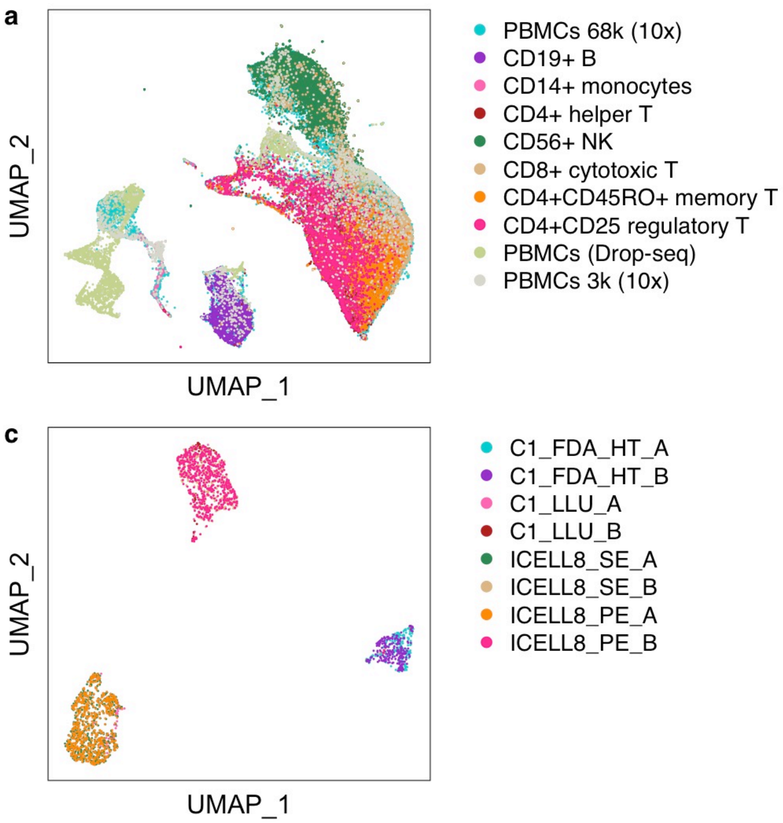


**Supplementary Figure 24. Scanorama batch correction using 10X and/or non-10x scRNA-seq datasets.**

(**a-b**) UMAP projections of PBMC datasets (10 subsets out of 26 datasets from Hie et al. paper, *Nature Biotechnology* 37, 2019) including two different technologies (10X Genomics and Drop-seq) before (**a**) or after (**b**) batch-effect correction. Datasets are displayed with two different colors (**b**) by technology (10X, blue color; Drop-seq, purple color). (**c-d**) UMAP projections of non-10X datasets (8 subsets out of 20 datasets from our own study) including three different non-10X technologies before **(c)** or after **(d)** batch-effect correction. Datasets are displayed with three different colors (**d**) by technology (C1_FDA_HT, blue color; C1, purple color; ICELL8, pink color). The PBMC datasets are downloaded from http://scanorama.csail.mit.edu/data_light.tar.gz. Our eight datasets were preprocessed using the featureCounts pipeline and batch-effect correction was performed using Scanorama V1.4.

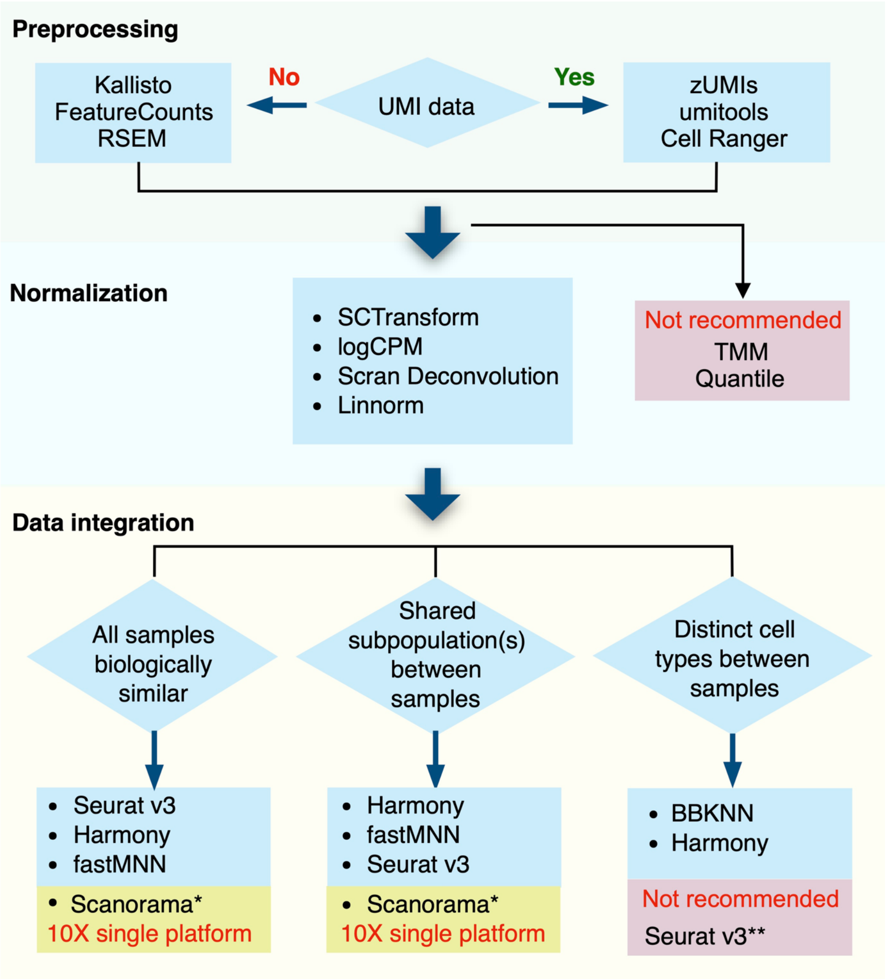


**Supplementary Figure 25. Best practice recommendations for single-cell RNA-seq analysis.**

*The current version of Scanorama did not correct batch effects for data from multiple platforms; however, it worked well when only 10X Genomics data were analyzed.

**Seurat v3 was suitable for the biologically similar samples, but over-corrected batch effects and misclassified cell types if large fractions of distinct cell types were present in different batches.
